# Physicochemical Properties of Enhanced and Recovered Venezuelan Crude Oil Treated with Biocompatible Pharmaceutical Formulation: Simulation of The Role of Water and Ionic Content

**DOI:** 10.1101/2025.07.26.666956

**Authors:** José Gregorio Colina-Moreno, Efrén de Jesús Andrades, Hilda Cristina Grassi, Lucila Estefanía Oquendo-García, María Alejandra Fernández-Ramírez, Anyi Yannire López Ramírez, Erick Alejandro Pacheco-Collazo, y Víctor Juan Andrades-Grassi

**Author notes:** This work is a Posthumous Tribute to Professor Hilda Cristina Grassi. Corresponding autor.

## Abstract

In this work, the hydrogen bonding interaction of water and ethanol with Polysorbate 80 (P-80, Tween 80) is simulated, in relation to its role in oil upgrading and recovery, using real theoretical and experimental studies. The theoretical physicochemical results obtained were compared with other experimental results derived from the treatment of improved and recovered oil after mixing Biocompatible Pharmaceutical Formulations (BPF) of these three compounds and oil produced in Venezuela. The Spartan 16 program was used to build the molecular structures and simulate the interactions with the semi-empirical PM3 method, to compare them with two real experiments. These two experiments were performed as follows: in the first, the mixture consists of a surfactant, (Z)-Sorbitan mono-9-octadecenoate poly(oxy-1,2-ethanediyl) or polyoxoethylene (20) sorbitan mono-oleate (Polysorbate 80), which is mixed with the hydrocarbon, and then completely fluidized with the addition of ethyl alcohol or ethanol.Water and ions (<100 ppm) present in this experiment are contained in oil and ethanol as contaminants (approximately 1.73-1.30% of total water). In the second experiment, the BPF ratios were 10:5:0.1:0.01 for oil:water with 17,800 ppm NaCl:P-80:ethanol (approximately 33.09% of total water and 1.78% w/v NaCl). The simulations were performed taking into account both real experiments. The simulated interactive products were characterized by the following parameters: energy, minimum conformational energy, solvation energy, aqueous energy, HOMO energy, LUMO energy, ΔE, dipole moment, polarizability, electrostatic potential map (EPM) and electron density.A theoretical parametric relationship (***sio Index***, “stability in oil Index”) is developed to evaluate the stability of the molecular forms generated within the hydrocarbon. The following theoretical experiments were performed: First, successive interactions of water with P-80 from position 1 to position 21. Second, successive interactions from position 1 to position 21 of P-80 of water and ethanol, 1 to 1. Third, successive interactions with P-80 of 21 ethanol molecules on the final product of the first experiment. These theoretical results were compared with some experimental results obtained using Oil Mixtures. The results obtained and compared in both experimental series (theoretical and real) allow to distinguish the role of P-80, Ethanol, Water and ions for the improvement and recovery of the treated oil. It is concluded that the formulation that does not contain added water and sodium chloride is supposed to use the contaminating water present in the petroleum and the water contained in the ethanol (<5%).This formulation has very little effect on the resulting Density of the improved Oil, but it greatly decreases the resulting Viscosity in the Crude Oil. The formulation that contains more than 30% added Water decreases the Density while the Viscosity of the Crude Oil responds proportionally to the Water retained in it. This difference in the results could be interpreted as a different proportion and chemical composition as a consequence of a different structural relationship between these components and a different conformation of the nanoparticle generated within the Oil, in each case. In this sense, the hypothesis is established that there would be an intermediate molecular form that could improve both Viscosity and Density.

## Introduction

Water is the liquid that dissolves the most substances (universal solvent). This property is due to its ability to form hydrogen bonds and additional interactions with other substances, since the latter dissolve with the polar molecules of water. One problem addressed by this work is the presence of water (permanent dipole) and ions with negative and/or positive charges, such as Sodium Chloride in Petroleum. The association of some polar, but non-ionic surfactants such as Polysorbate 80 (P-80) with other organic molecules such as Choline Chloride [1],and in our case, Ethanol [2] can solve some problems such as desalination on the one hand and the presence of water (permanent dipole) and ions with negative and/or positive charges, such as Sodium Chloride in Petroleum, respectively, through the interaction of Hydrogen bonds, Van der Waals forces and others.

The interaction between a permanent dipole and an induced dipole explains how hydrocarbons, such as oil (less polar molecules) relate to water (polar molecule):the permanent dipole of the polar molecule deforms the charge cloud of the less polar molecule, generating an induced dipole in the latter, depending on the Dipole Moment (DPM) and the Polarizability (P).Finally, an interaction occurs between the permanent dipole of the polar molecule and the induced dipole in the less polar molecule [3].

Petroleum hydrocarbons can hardly be affected by Water to form an induced dipole. This is why the introduction of other organic molecules such as P-80 and Ethanol in Biocompatible Pharmaceutical Formulations (BPF) can affect this situation due to their DPM, P and other properties that can improve the physicochemical relationship of Water and ions with hydrocarbons.This work attempts to highlight this situation by evaluating these physicochemical parameters, the potential stability of the molecules within the oil and, more importantly, the improvement and recovery of the hydrocarbon in terms of its viscosity and density [2, 4].The induction in a molecule of a dipole that previously did not exist can cause the expulsion of that molecule from the Petroleum.In this sense, the Hydrocarbon would become a “Selector” of those molecules that meet an MDP and a P suitable for their stable permanence within it.In this work, the theory and the experimental part that justify these concepts for Venezuelan crude oil are developed.

In summary, the hydrophobic interaction between water and petroleum is based on the inability of less polar hydrocarbon substances to form hydrogen bonds with water molecules [5].This problem has been partially solved in this work by the interaction of P-80 associated with Ethanol, in the presence of Petroleum containing Water and ions.

From previous work [2] it is concluded that this BPF is minimalist in terms of the proportion of water required for viscosity improvement. In this first formulation,the salts present in the mixture, including NaCl, are exclusively those present in the hydrocarbon. According to the results of this work, the structures generated by the interaction of hydrogen bonds of water and ethanol with P-80 that resulted in an improvement of viscosity in that work, should be stabilized as oligomers with a relatively tubular or prolate structure (“Prolate ellipsoid of revolution”) [6], If compared to the structures proposed for the P-80 by Aizawa 2009 [7] which have both a tubular and spherical structure, the latter would be oblate (“Oblate ellipsoid of revolution”) [6].*the* BPF *of the second real experiment of the present work [4] would lead to this last type of structure with the consequent improvement of the density of the Petroleum. In this work these hypotheses are confirmed,and a new one is proposed: There could be a new molecular formula intermediate between the flat and spherical ones, which would improve both Viscosity and Density*.

## Materials and methods

The following theoretical experiments were performed: First, successive interactions of water with P-80 from position 1 to position 21. Second, successive interactions from position 1 to position 21 of P-80 of water and ethanol, 1 to 1.Third, successive interactions one by one up to 21 ethanol molecules with P-80 on the final product of the first experiment.

These theoretical results were compared with some experimental results obtained using BPF Blends with Petroleum.In this work, an empirical relationship was developed between the difference in HOMO and LUMO Energy (ΔE), with the DPM and P, which would indicate the value of the Stability Index (sio Index “stability in oil index”) of each molecule in the crude oil,as follows (see Table 5):

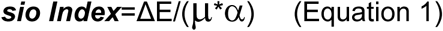

***sio Index*** = Stability; ΔE = Energy Difference, E_HOMO_-E_LUMO_;μ = Dipole Moment; α= Polarizability;

In this equational relationship the numerator highlights the potential reactivity of each Molecular Form that derives from the difference in HOMO-LUMO Energy [10].This reactivity would be important to achieve a stable interaction, for example, between two complementary monomers of each Molecular Form, to form a dimer and further an Oligomer [see Figure 1 (B, C and D) below]. On the other hand, the denominator would highlight the role of the greater or lesser potential Polar or ionic character of each Molecular Form and with it the greater or lesser potential tendency for it to be rejected by the Petroleum.To facilitate visualization and handling, sio Index values are shown in texts, tables and graphs as values between 1 and 100 multiplied by 10^3^ with two significant decimal places.However, exact results are available with decimal-only values up to 6 significant figures.

**Figure 1(A).**
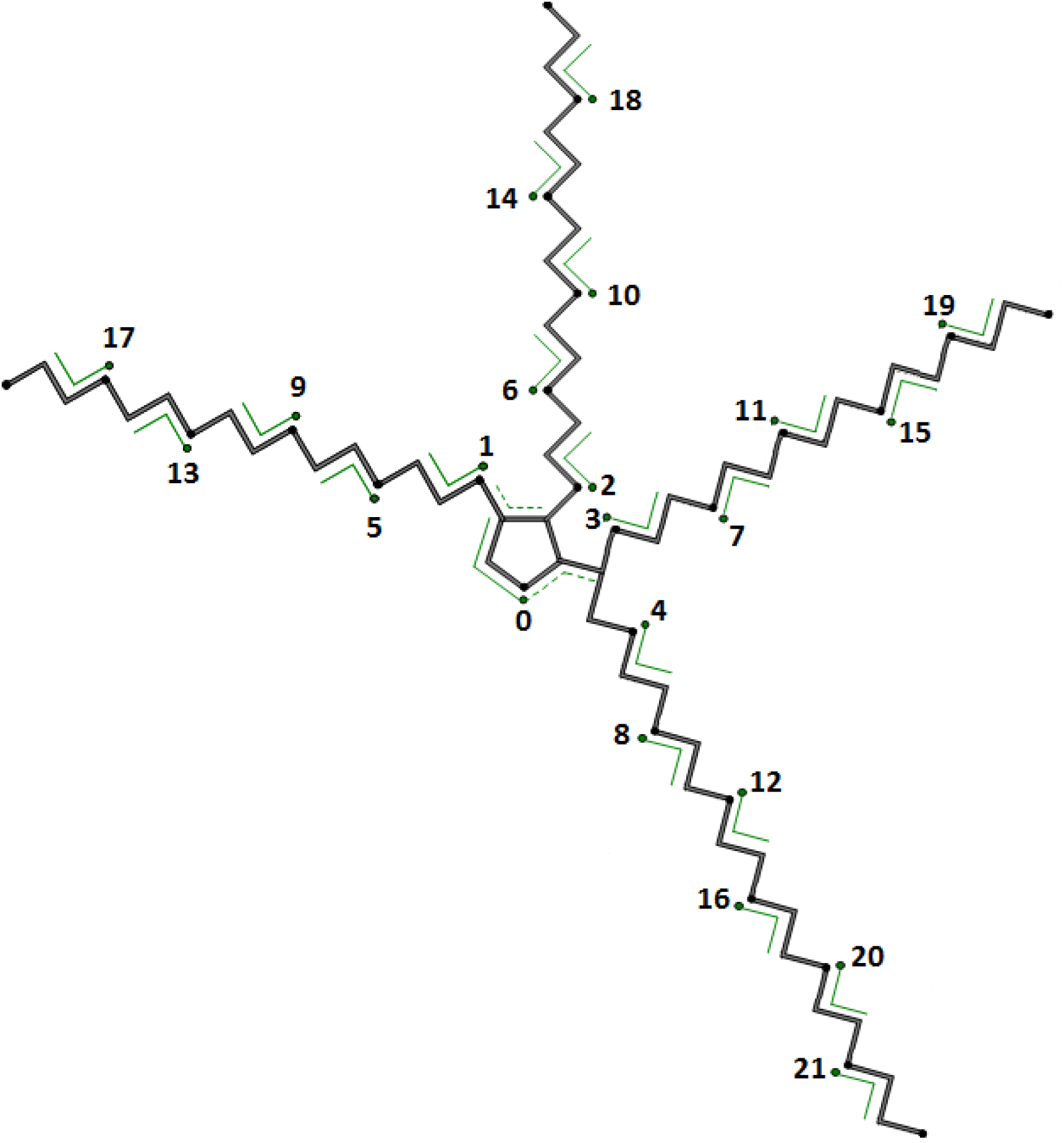
Candidate positions for establishing hydrogen bonds with water and/or ethanol in a P-80 molecule with the 21 ethoxides and without the C-17 hydrocarbon extension.

The structure of the P-80 molecule is shown in the design of Figure 1 (A). In this Figure the zero position is highlighted which is the Oxygen of the Carbonyl Carbon of the Sorbitol ring, this position is not considered in some cases, because the potential Hydrogen bond in it does not participate in the structure of the Ethoxides.In most figures in the supplementary material an arrow is used to indicate this position. The stability of the P-80 ring, is ensured by the hydrogen bond between water and the oxygen that closes the ring, without altering the structure of P-80 [8]. The numbering of the positions for successive interaction of water was chosen, or water and ethanol using a thermodynamic structural criterion [9] which defines that the most stable interactions are closer to the central structure of sorbitol and from there towards the periphery following an order of proximity to it.This design influences the outcome of both individual and successive interactions, as will be seen later in results. Although the interaction with water (Simulation Series 1) and with water and ethanol simultaneously (Simulation Series 2 and Series 3) and the oxygen of sorbitol were simulated, these interactions are reported separately, since the relevant interactions are expected to be with the oxygens of the ethoxides. This consideration is made because the sugar ring would tend not to be destabilized by this type of interactions [8].

The Spartan 16 program was used to construct the molecular structures and simulate the interactions with the semi-empirical PM3 method. The simulated interactive products were characterized by the following parameters: Energy, Minimum Conformational Energy, Solvation Energy, Aqueous Energy, HOMO Energy, LUMO Energy, ΔE, Dipole Moment, Polarizability, Electrostatic Potential Map (EPM) and Electron Density.

The results of the theoretical experiments were compared with two real experiments trying to distinguish the effect of the more hydrophilic polar and/or ionic components such as water and sodium chloride. In the first experiment [2] two formulations were used: Mixture 1: P-80, two parts; Petroleum, one part (dissolved ionic solids <100 ppm; <0.01%); Ethanol, one part (1.30% total water). Mixture 2: P-80, one part; Petroleum, one part (dissolved ionic solids <100 ppm; <0.01%); Ethanol, one part (1.73% total water).

In the second experiment [4] the following formulation was used: P-80, 0.1 part; Petroleum, 10 parts; Ethanol, 0.01 part; Water containing 17,800 ppm NaCl (1.78%), 5 parts.The viscosity and density, among other parameters, were measured in both experiments of the upgraded and recovered petroleum [2, 4].As will be seen in the results, the mass ratio in the formulations between P-80 and Water could be defining the improvement of the Petroleum in its viscosity and density.

This work attempts to compare the results of the simulation with the results of real experiments, emphasizing the generated structures and some physicochemical parameters and, on the other hand, the real contribution of the experimental results to the improvement and recovery of petroleum extracted in Venezuela. In this way, the molecular structures generated by the simulation that would possibly induce petroleum upgrading and recovery could be selected, according to the viscosity and density results obtained in real experiments.

## Results and discussion

In the first theoretical experiment (Series 1), Figures 2 (B) to 23 (B), which show the result of the successive interactions of water with P-80 from position zero to position 21. The results obtained show that the molecular form of Figure 5 (B) presents a minimum Dipole Moment (DPM) of 3.09 Debye in this experiment (see Table 1). However, the Polarizability (P) of all the molecular forms generated in this first experiment exceed the value of154,71 C m^2^/V. The DPM value of 3.09 would imply the molecular form interacting with three water molecules would be in better condition than P-80 in its original form to enter the hydrophobic phase of the Petroleum [see EPM in Figure 24 (B)]. However, the second value (P) would imply that all molecular forms in Table 1,they would not have the appropriate conditions to remain within the crude in the event that the concentration of polar or ionic molecules present in the aqueous phase (for example, such as Sodium Chloride) was very high.This would be caused by the induced dipole-dipole interactive phenomenon. That is to say, the molecular form of Figure 5 (B), although it has a low DPM, would be subject to a change in polarity induced by the salts present in the aqueous phase of the petroleum and could be expelled from it. This means that, in the case of this first simulation, all the molecular forms that have the possibility of entering the crude oil would depend, for their permanence in it, on the concentration of ions present in the Petroleum. The result of this first simulation justifies its comparison with the two real experiments: low concentration of Sodium Chloride and high concentration of Sodium Chloride, first and second real experiments, respectively (see Materials and Methods).

Trying to modify the Physicochemical parameters obtained in the first simulation with P-80, a second series was carried out in which the successive interactions from position 0 to 21 were carried out with a water molecule and an Ethanol molecule simultaneously.The first important change is obtained in the DPM with 2.52 and P of 176.85 in the interaction of 4 Water molecules and 4 Ethanol molecules simultaneously [see Figure 6 (C) and EPM Figures 25 (B) and 25 (C)], unlike the first series which was obtained with 3 Water molecules (DPM= 3.09). Moreover, the value of P drops drastically to 38.83 by interacting with 11 Water and Ethanol molecules simultaneously [see EPM of the series of Figures 26, 27, 28, 29 and 30 all (B) and (C)].This phenomenon will be explained later in this work (see Table 1, 2 and 3. This phenomenon speaks of the stability of molecular forms). In this series, The ideal conformational form that can enter the crude oil and remain in it would be the one derived from the interaction of P-80 with 13 Water molecules and 13 Ethanol molecules simultaneously [Figure 15 (C), DPM= 4.15 and P=38.48].As seen in Figures 1 (B, C and D) (see below), it is possible for Dimerization or Oligomerization to occur with several of these molecules by coupling the “groove” on one side of one molecule with the “hump” of the other molecule.The charge densities of these two molecular forms are complementary. This P-80 molecule could withstand relatively high ionic concentrations to remain within the crude oil. In this work, the hypothesis is established that this series of intermediate molecular forms could improve the viscosity and density of the crude oil at the same time.

Trying to highlight the role of water and Sodium Chloride, a third series of simulations was tested by adding to the last molecular form generated in the first series, see Figure 23 (B), successively from 1 to 21 ethanol molecules occupying positions 1 to 21, which were already interacting with water. The result obtained is shown in Figures 2 (D) to 23 (D) and in Table 3. The results obtained show that the molecular form in Figure 7 (D) presents a minimum DPM of 2.90 in this experiment (see Table 3). However, the Polarizability (P) of this molecular form generated in this case is 209.71. This last value would not allow this molecular form to remain within the crude oil at relatively high ion concentrations. The P values drop dramatically when 7 Ethanol molecules interact, this phenomenon will be explained later in this work (see Table 1, 2 and 3. Again this phenomenon speaks of the stability of the molecular forms).When it interacts with 9 Ethanol molecules, see Figure 10 (D), an DPM of 3.09 and a P value of 38.86 are obtained. This molecular form would be able to enter the crude oil and remain there, even at relatively high ionic concentrations. As can be seen in detail in Figure 10 (D) there is a succession of “layers”, the first of ethoxides, the second of water and the third of ethanols, the latter covering the molecule on the outside, It is possible that this molecular form expresses the ideal situation for the DPM and P to be necessary to enter and remain in the oil and improve its density.

From Figures 2 (B) to 23 (B) it is observed that successive interactions with water produce a conformational change from linear to spherical, always preserving the exposed hydrocarbon extension and possibly available to interact with petroleum. As will be seen later, the linear or flat forms of this molecule would contribute to improving the viscosity of the crude oil.

In this work we try to compare the simulation results with the results of real experiments, emphasizing the generated structures and some physicochemical parameters andon the other hand in the real contribution of the experimental results in the improvement and recovery of petroleum extracted in Venezuela. In this way,molecular structures generated by the simulation could be selected that would possibly induce petroleum upgrading and recovery, according to the viscosity and density results obtained in the real experiments.

In this sense we have two bibliographical references [2, 4], from which part of the relevant data can be extracted to compare with series 1, 2 and 3. In the first real experiment [2] an attempt was made to improve and recover the Oil with two formulations with excess Oil, P-80 and Ethanol and with a marked deficit of Water and ions. In the second real experiment [4], an attempt was made to improve and recover the Oil with a formulation with excess Oil and with excess Water and ions and a deficit of P-80 and Ethanol. The result is clear: in the first formulations the Viscosity was improved and not the Density, and in the second formulation it tends to be the opposite. The same thing happens in this last case with other linear n-alcohols from C-1 to C-8, with which a better improvement and recovery result is obtained in Petroleum Density for even alcohols and worse for odd ones. Proportional results are also obtained from the Viscosity of the Oil in relation to the Water retained in it: the greater the amount of Water retained in the Petroleum, depending on the alcohol used, higher the Viscosity and lower the Elasticity of the recovered Petroleum [4]. The latter coincides with the results obtained in the first experiment: in this case, the Viscosity of the upgraded and recovered Oil is decreased in the absence of Water and added ions, while the Density remains unchanged.

The result of series 1 aligns with the previous step of adding only P-80 to the Petroleum which occurs as described in Mixtures 1 and 2 of the first real experiment [2]. In this case, as described, P-80 mixes well with the oil and the accompanying contaminant water, but does not achieve its upgrading and total recovery. This can be observed in Figure 5(B) and Table 1, (P-80 semiempirical-3 molH2O (P-80 + 3 mol. Water; Pos. 1-3). This result justifies the simulation of series 1 in that there is an association of the few water molecules contained in the crude oil with the P-80 within the oil,in addition to justifying the consideration that under normal well conditions or in conditions of already produced petroleum, we have two methods of evaluating improvement and recovery, mainly of viscosity, one simulated and another real under water deficit conditions [2].

The result of series 2 aligns with the subsequent step of adding Ethanol to the result obtained in the previous paragraph, which occurs in Mixtures 1 and 2 of the first real experiment [2]. This can be observed in Figure 6 (C) and Table 2, (P-80 semiempirical-4 molH2O-4 mol-ethanol (P-80 + 4 mol. Water + 4 mol. Ethanol; Pos. 1-4). This result justifies the simulation of series 2 in that there is an association of the water molecules contained in crude oil and ethanol with P-80 within the Petroleum, in addition to justifying the consideration that under normal well conditions or in conditions where the Petroleum has already been produced, we have two methods of evaluating improvement and recovery using ethanol, one simulated and one real.

An even more interesting result is observed in this series 2 in the molecular shapes ranging from P-80: 11 Water molecules: 11 Ethanol molecules, to P-80: 15 Water molecules: 15 Ethanol molecules, with an optimal result at P-80: 13 Water molecules: 13 Ethanol molecules. This can be observed in Figure 15(C) and Table 2, (P-80 semiempirical-13 molH2O-13 mol-ethanol (P-80+13 mol. Water+13 mol. Ethanol; Pos. 1-13). This molecular form, which was discussed at the beginning of the results and discussion section, is also shown in Figure 1 (B, C and D). The existence of this molecular structure, its Dimer and its possible Oligomer allows us to hypothesize that it is an intermediate form both from the point of view of the association of the number of Water and Ethanol molecules and the possible improvement of the Density and Viscosity of the Oil at the same time.

**Figure 1(B, C and D).**
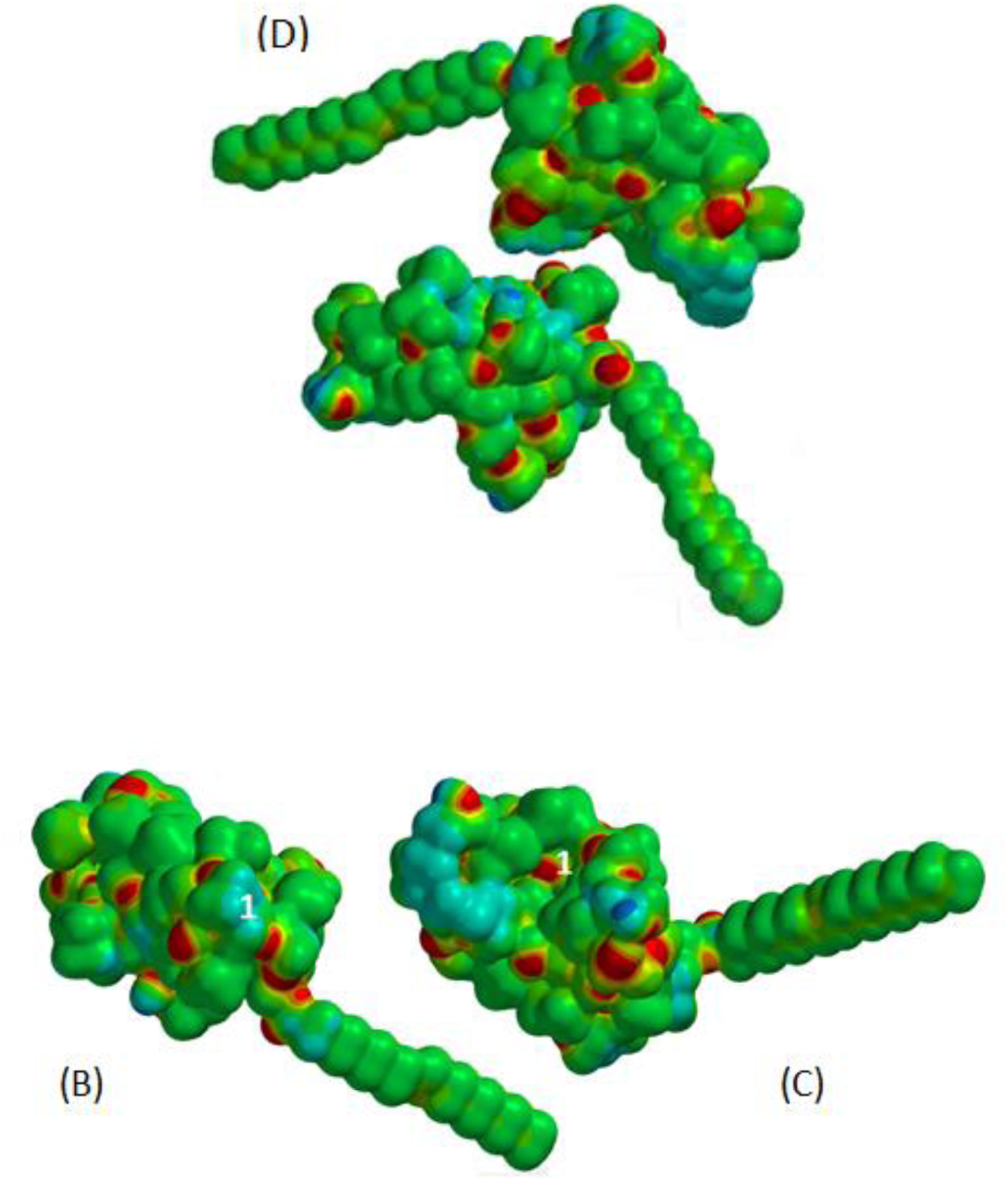
Electrostatic Potential Maps (EPM): On the top left (B): the ionic groups with positive charge density (1 blue) of the “hump” of the P-80:13 water:13 Ethanol molecule are shown and on the top right (C): the ionic groups with negative charge density (1 red) of the “groove” of the P-80:13 water:13 Ethanol molecule are shown. Both are coupled to form 1 Dimer shown in (D).

The result of series 3 is in line with the second real experiment (see Table 5) [4], which is performed with a deficit of P-80 and an excess of water and ions. This can be observed in Figure 7 (D) and Table 3, ((0-5) + P-80 semiempirical-21 molH2O+ (0-5) mol ethanol [P-80 + 22 mol. Water; Pos. 0-21 + 6 mol. Ethanol; Pos. 0-5). On the other hand, it can also be observed in Figure 10(D) and Table 3, and also MPE in Figure 31(C) (0-8) + P-80 semiempirical-21 molH2O+ (0-8) mol ethanol [P-80 + 22 mol. Water; Pos. 0-21 + 9 mol. Ethanol; Pos. 0-8),a better result for the improvement and recovery of Oil under conditions of the following formulation applicable to both results (see materials and methods): P-80, 0.1 part: Oil, 10 parts: Ethanol, 0.01 part: Water containing 17,800 ppm NaCl (1.78%), 5 parts. These two results justify the simulation of series 3 in that there is an association of a total of 22 water molecules (added in excess) together with those contained in crude oil and ethanol with P-80 within the Petroleum,in addition to justifying the consideration that under normal well conditions or in conditions of already produced petroleum, we have two methods of evaluating improvement and recovery, mainly from the density, using ethanol, one simulated and one real under conditions of excess water.

### Table 1 Figure 2 (B) – 23 (B) (The arrow indicates the position of sorbitol)

It is observed that in the P-80 series only with water the Polarizability (P) value is high from the beginning to the end and increases gradually, while the DPM value oscillates between values slightly lower and slightly higher than the value of 4.06 of P-80 alone. This occurs up to 14 water molecules interacting with P-80 (2/3 of the total), after which the DPM values remain very high. This means that Polysorbate 80 mixed with Petroleum with low Water content and very low ionic concentrations (<100 ppm) can result in decreased Viscosity with improved oil recovery based on these results.

### Table 2, Figures 2 (C) – 23 (C) (The arrow indicates the position of sorbitol)

In series 2, in which the water and ethanol molecules are added 1 to 1, the value of P is higher up to 10 molecules of each. Again, in this case the criterion of the previous series applies, but here it is from 11 substitutions onwards. In this case, in addition to including Ethanol, the amount of Water may be greater, which could influence controlling moderate values for both Viscosity and Density.

As for the conditions for the induction of the formation of Oligomers [11] of molecular forms of P-80, homologous and heterologous, we will discuss them in this Series 2, in which Water and Ethanol are added 1 by 1. Before we get to the interactions that interest us for this possible outcome, we find the molecular form with P-80:4 Water molecules:4 Ethanol molecules (DPM of 2.52 and P of 176.85). This would only be maintained within crude oil with a low ionic level. The P value abruptly drops to 38.83 in P-80:11 water molecules:11 ethanol molecules and so on. Although only P-80:13 water molecules:13 ethanol molecules has a DPM (4.15), comparable to that of P-80 (4.06), we see in the molecular structures from 11 water molecules: 11 ethanol molecules, up to 15:15 (even earlier) a “groove” and a “hump” that would appear to be complementary (see Figure 1 (B, C and D) in the main text). This complementarity generates the hypothesis of a possible formation of self-assembled oligomers [11]. These self-assembled oligomers could generate a roller-like structure [12–13]. We also note that in this region the MDPM values are oscillatory. Oligomers could be formed equally in series 1 and 3, but their formation in series 2 is more evident.

**Figure 2.**
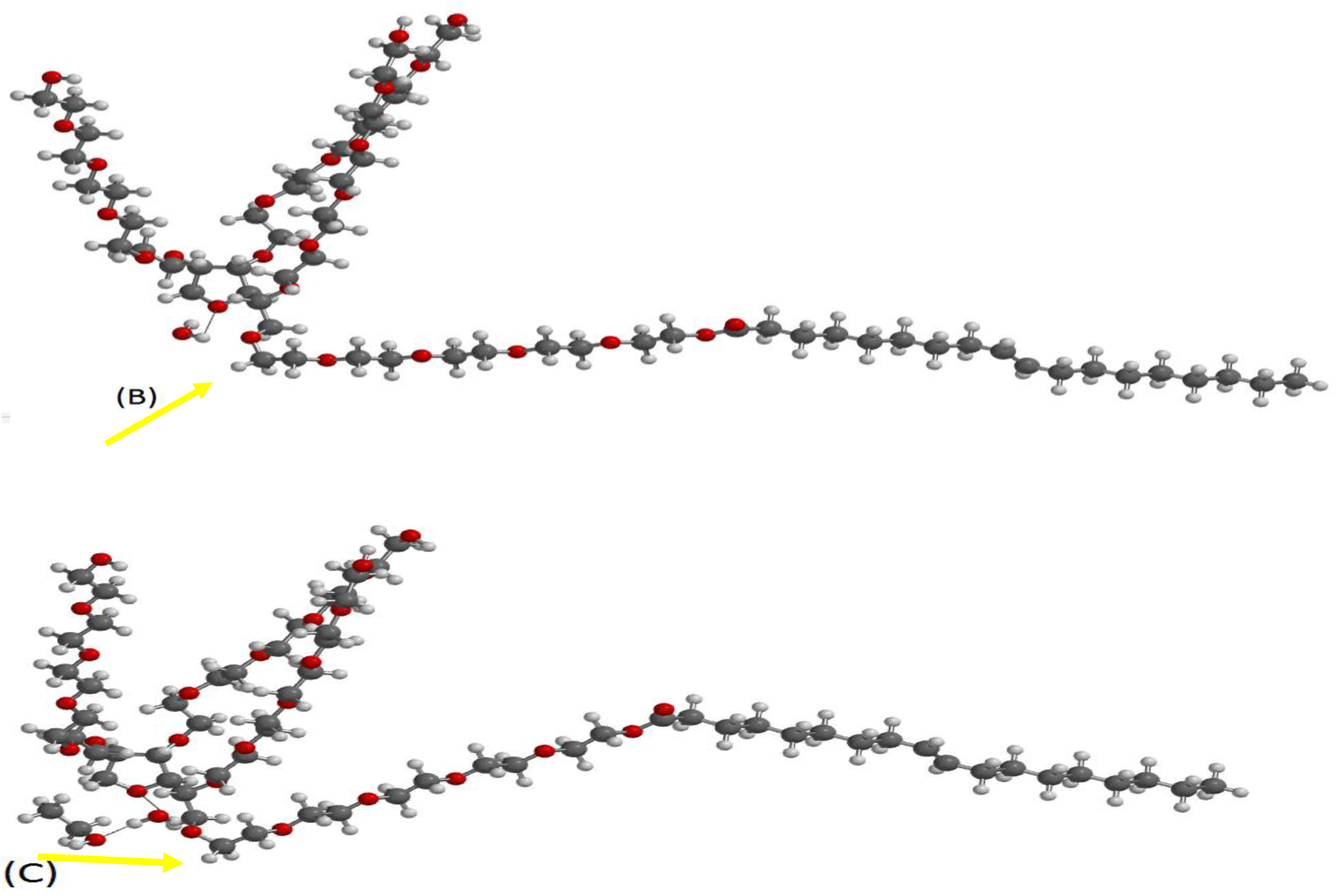
Comparison of (B) P-80 molecule interacting with 1 water molecule at position 0 (C) P-80 molecule interacting with 1 water molecule at position 0 and these in turn interacting with 1 ethanol molecule (CH_3_CH_2_OH).

**Figure 3.**
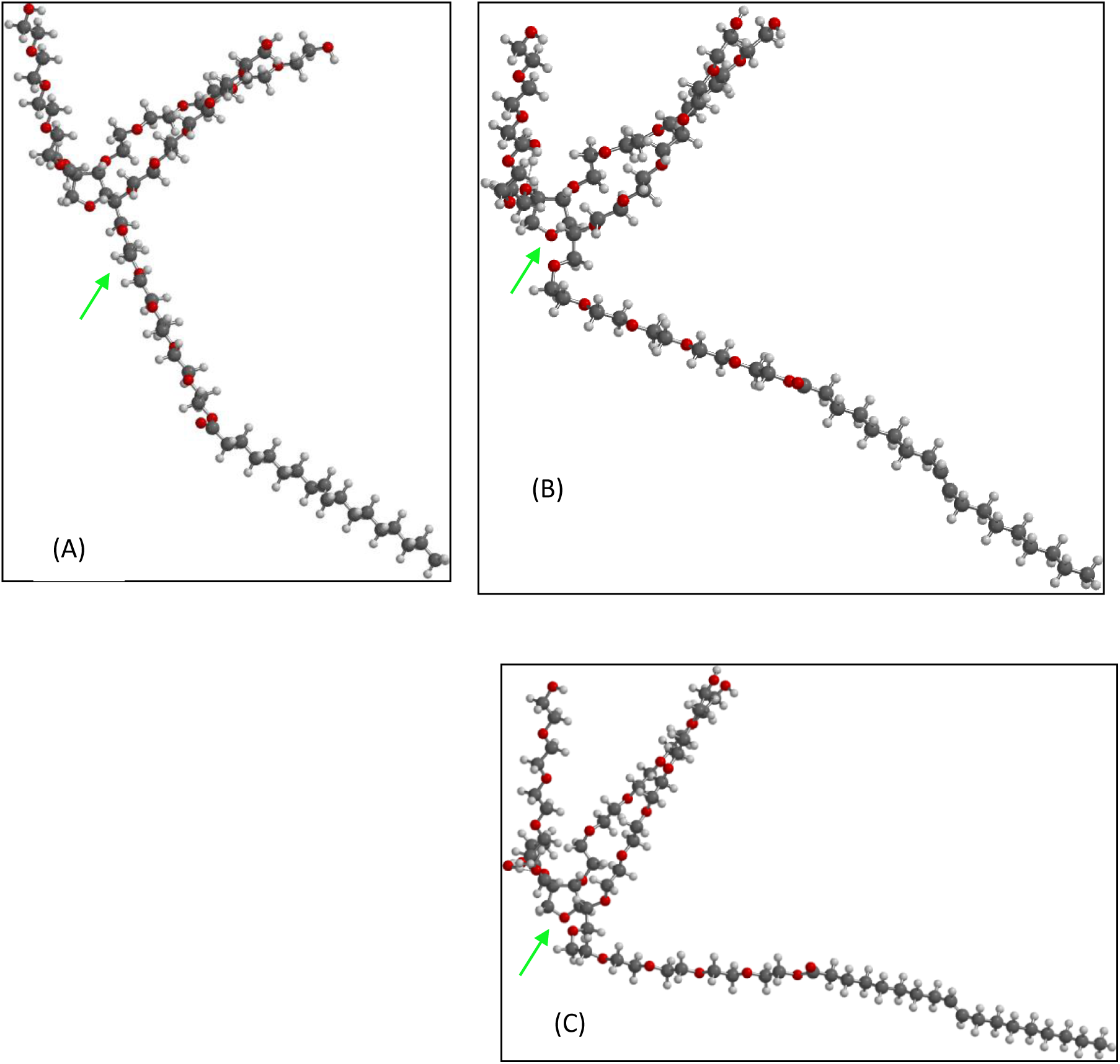
Comparison of (A) P-80 molecule without interaction; (B) P-80 molecule interacting with 1 water molecule at position 1; (C) P-80 molecule interacting with 1 water molecule at position 1 and this in turn interacting with 1 ethanol molecule (CH_3_CH_2_OH) at the same site.

**Figure 4.**
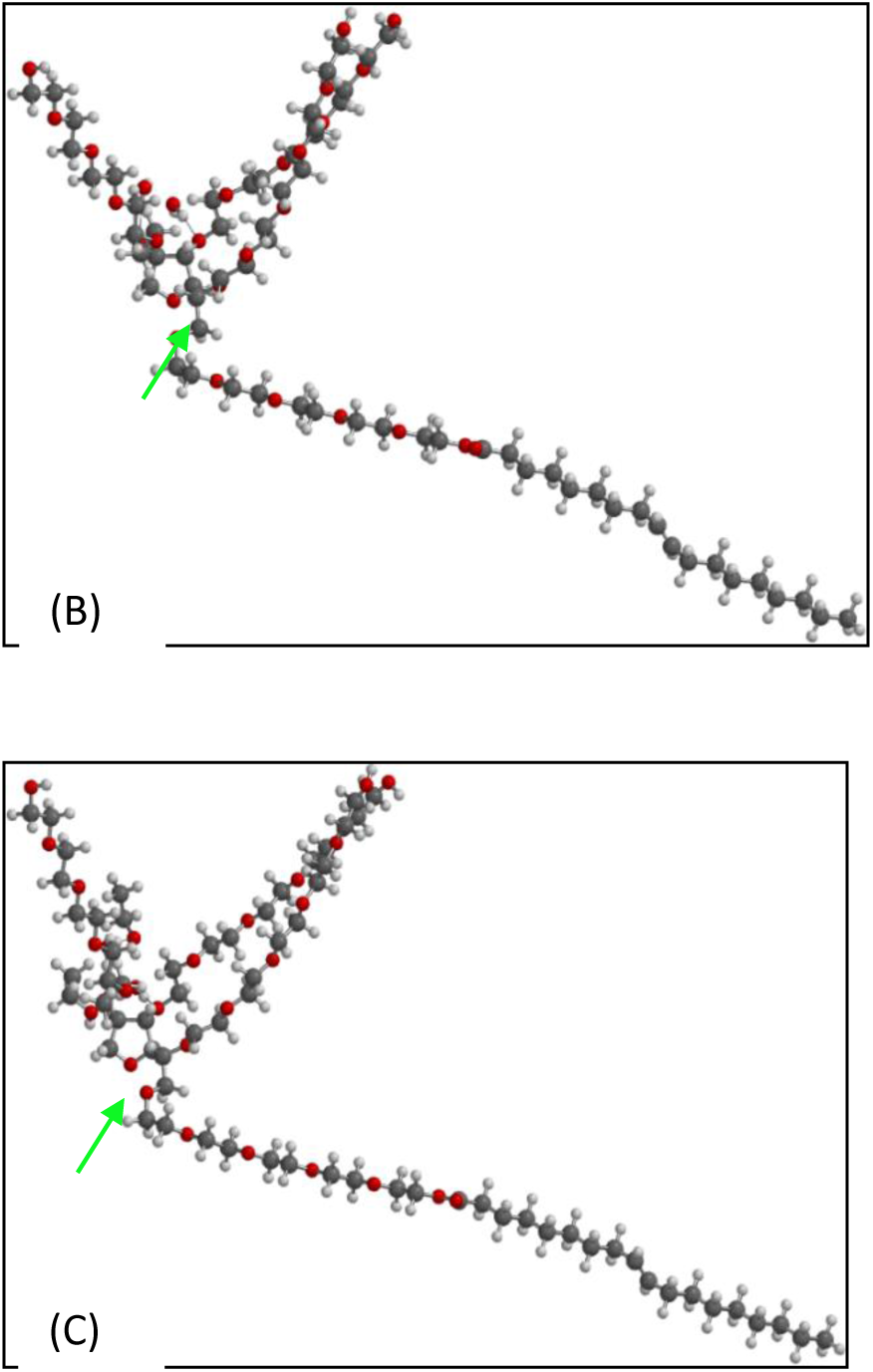
Comparison of (B) P-80 molecule interacting with 2 water molecules at positions 1-2; (C) P-80 molecule interacting with 2 water molecules at positions 1-2; and this in turn interacting with 2 ethanol (CH_3_CH_2_OH) molecules at the same site.

**Figure 5.**
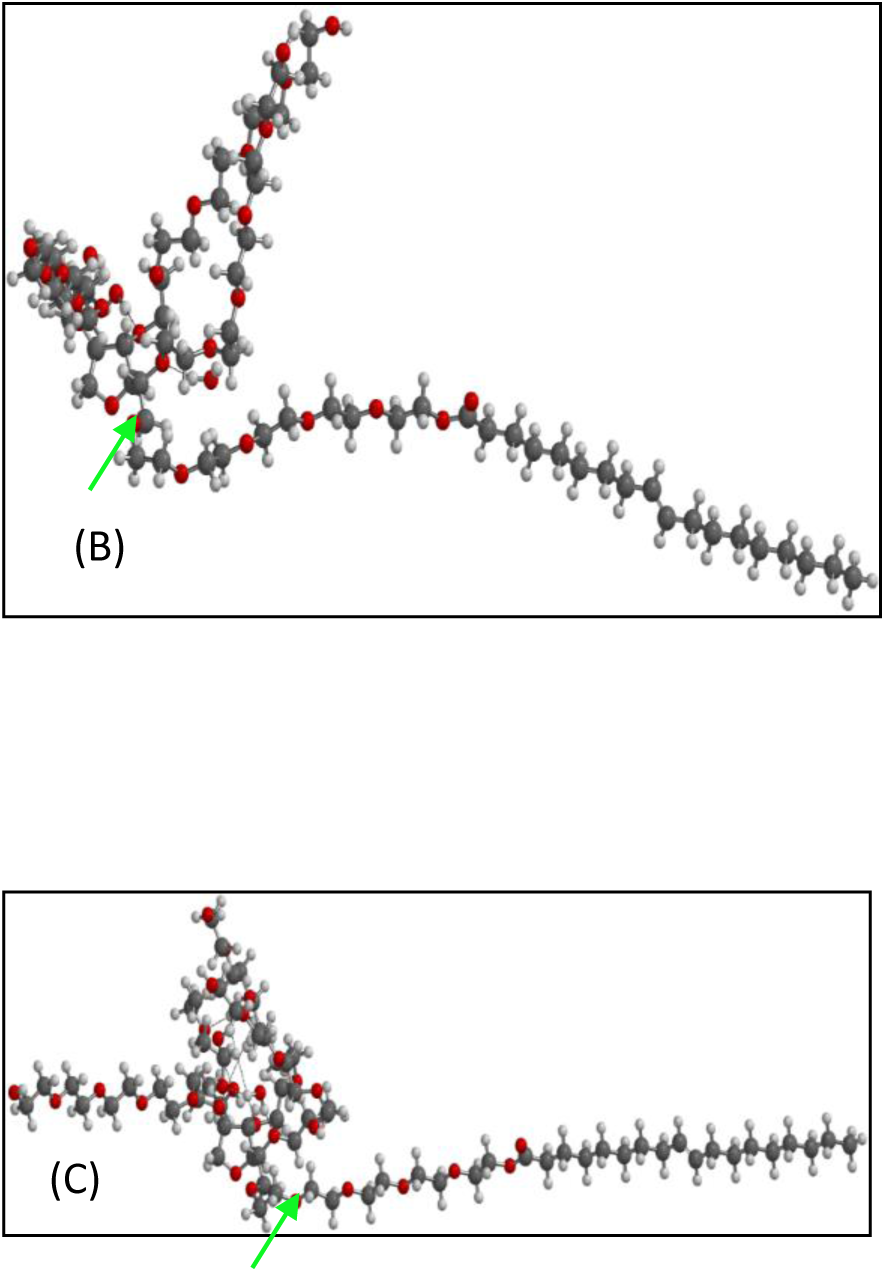
Comparison of (B) P-80 molecule interacting with 3 water molecules at positions 1-3; (C) P-80 molecule interacting with 3 water molecules at positions 1-3; and this in turn interacting with 3 ethanol (CH_3_CH_2_OH) molecules at the same site.

**Figure 6.**
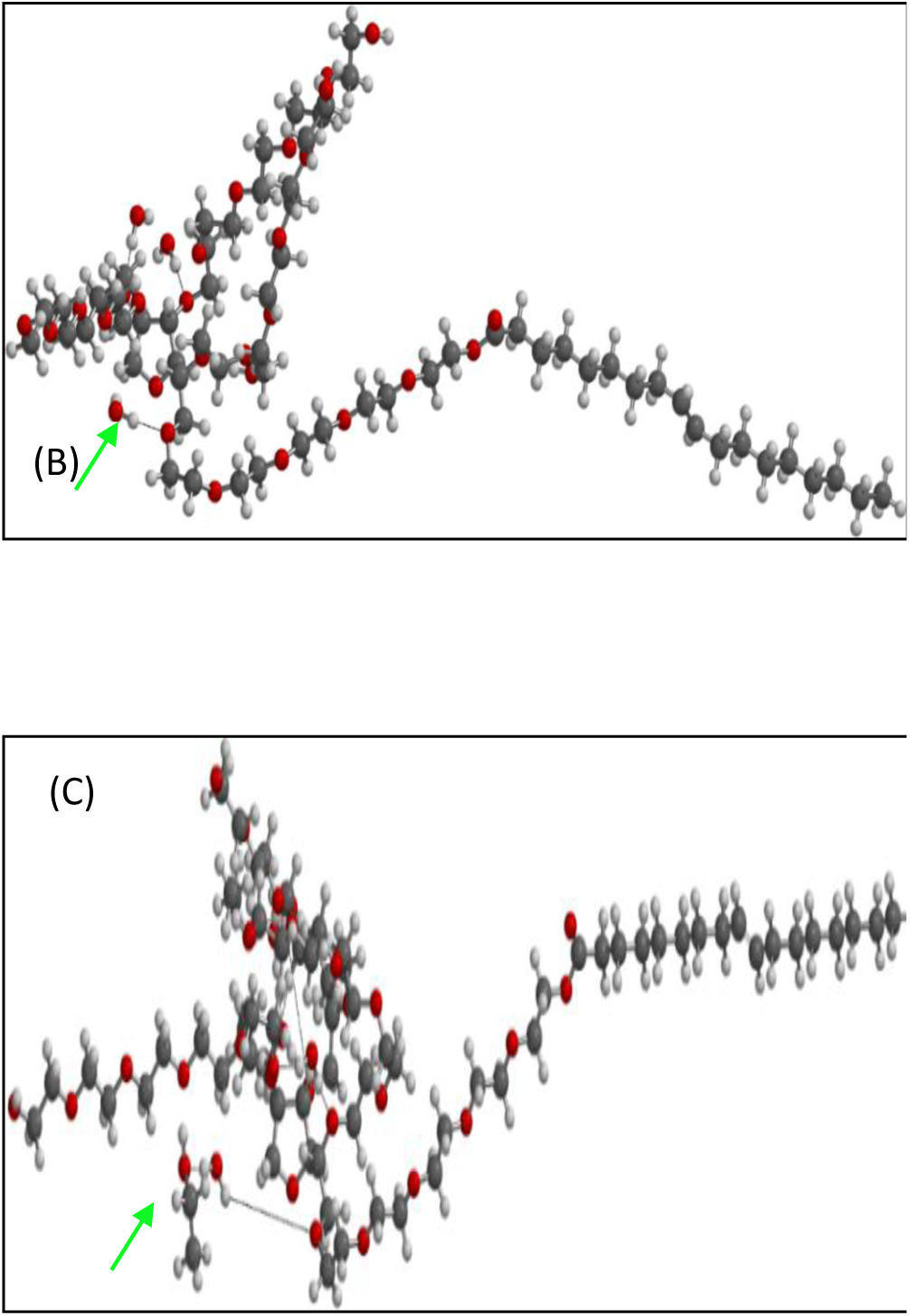
Comparison of (B) P-80 molecule interacting with 4 water molecules at positions 1-4; (C) P-80 molecule interacting with 4 water molecules at positions 1-4; and these in turn interacting with 4 ethanol molecules (CH_3_CH_2_OH).

**Figure 7.**
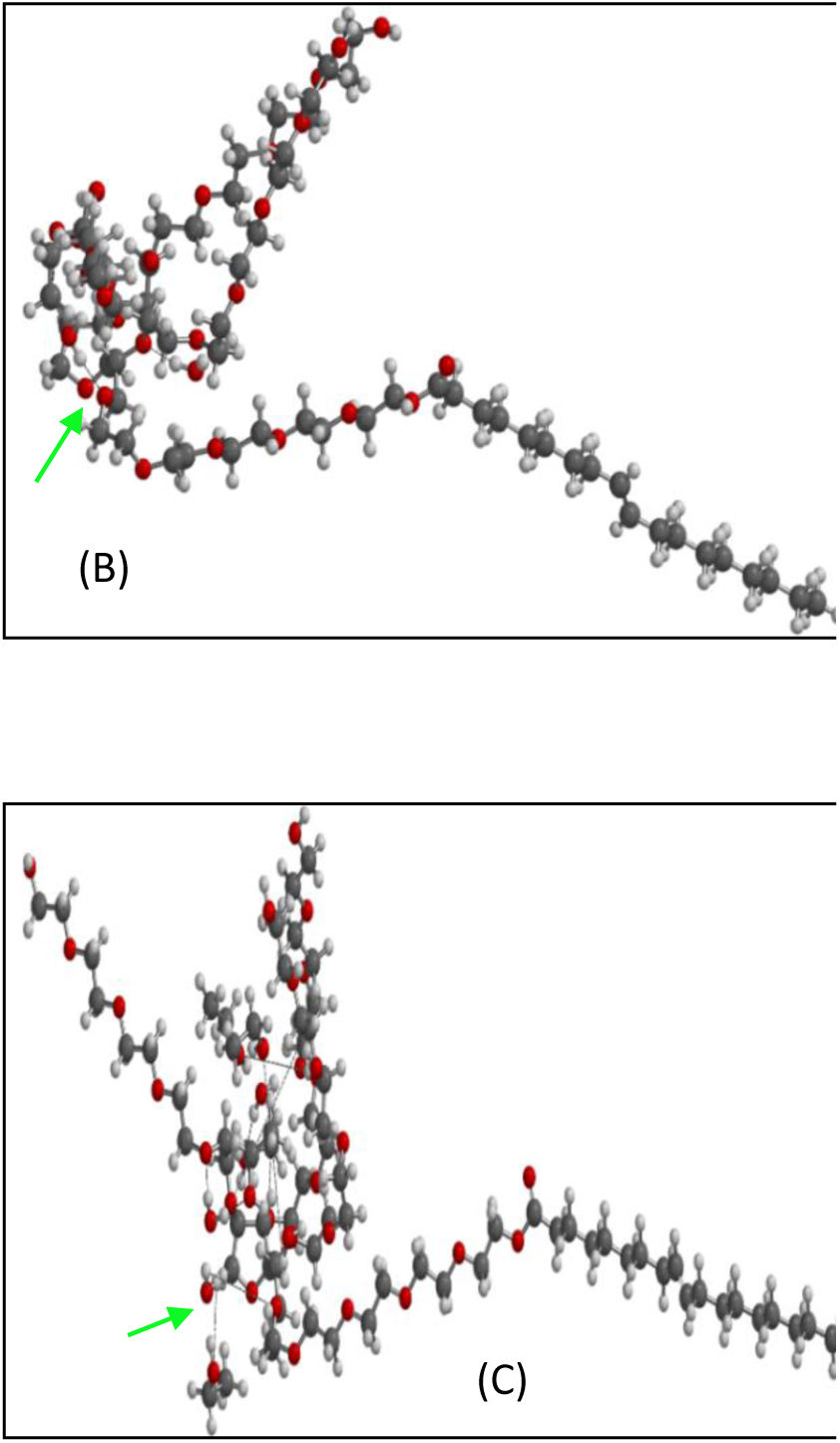
Comparison of (B) P-80 molecule interacting with 5 water molecules at positions 1-5; (C) P-80 molecule interacting with 5 water molecules at positions 1-5; and these in turn interacting with 5 ethanol molecules (CH_3_CH_2_OH).

**Figure 8.**
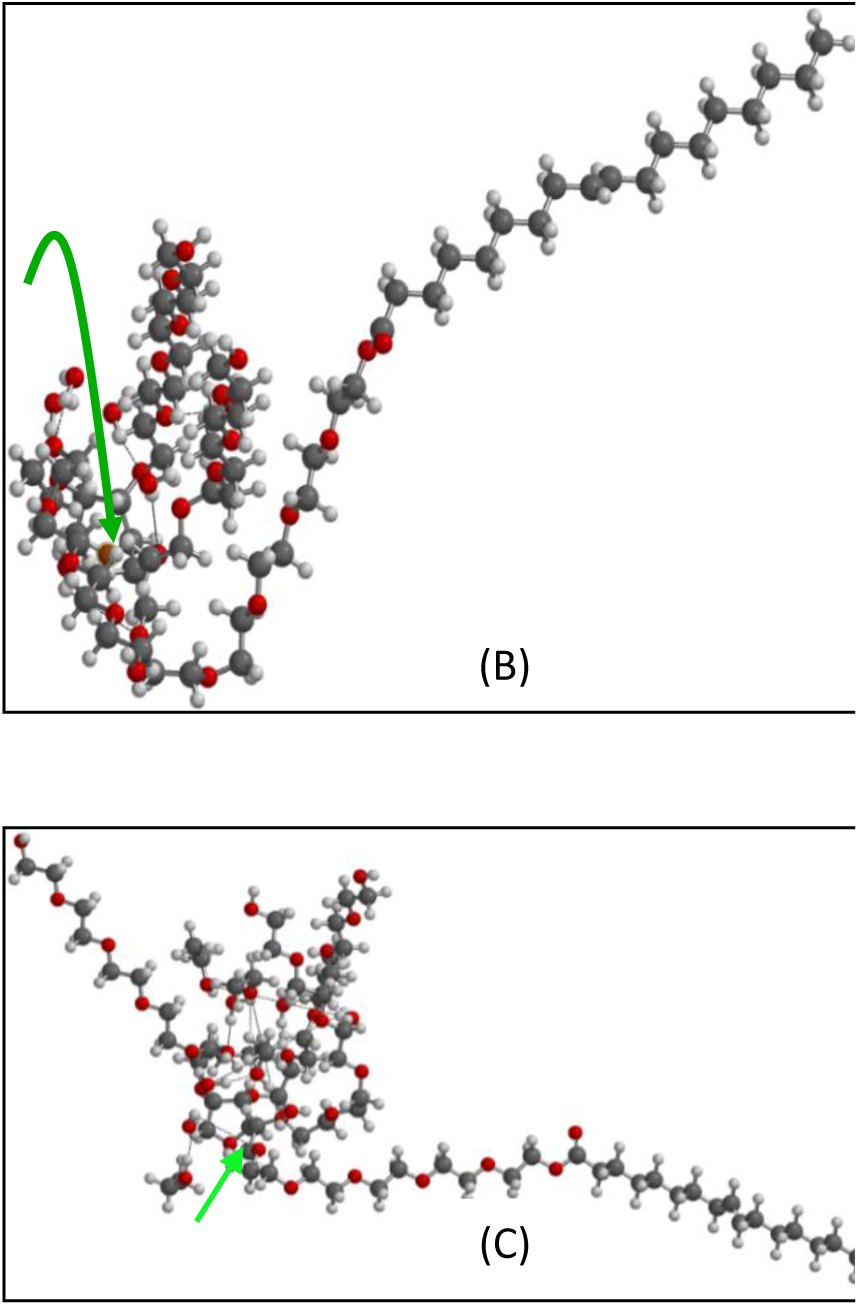
Comparison of (B) P-80 molecule interacting with 6 water molecules at positions 1-6; (C) P-80 molecule interacting with 6 water molecules at positions 1-6; and these in turn interacting with 6 ethanol molecules (CH_3_CH_2_OH).

**Figure 9.**
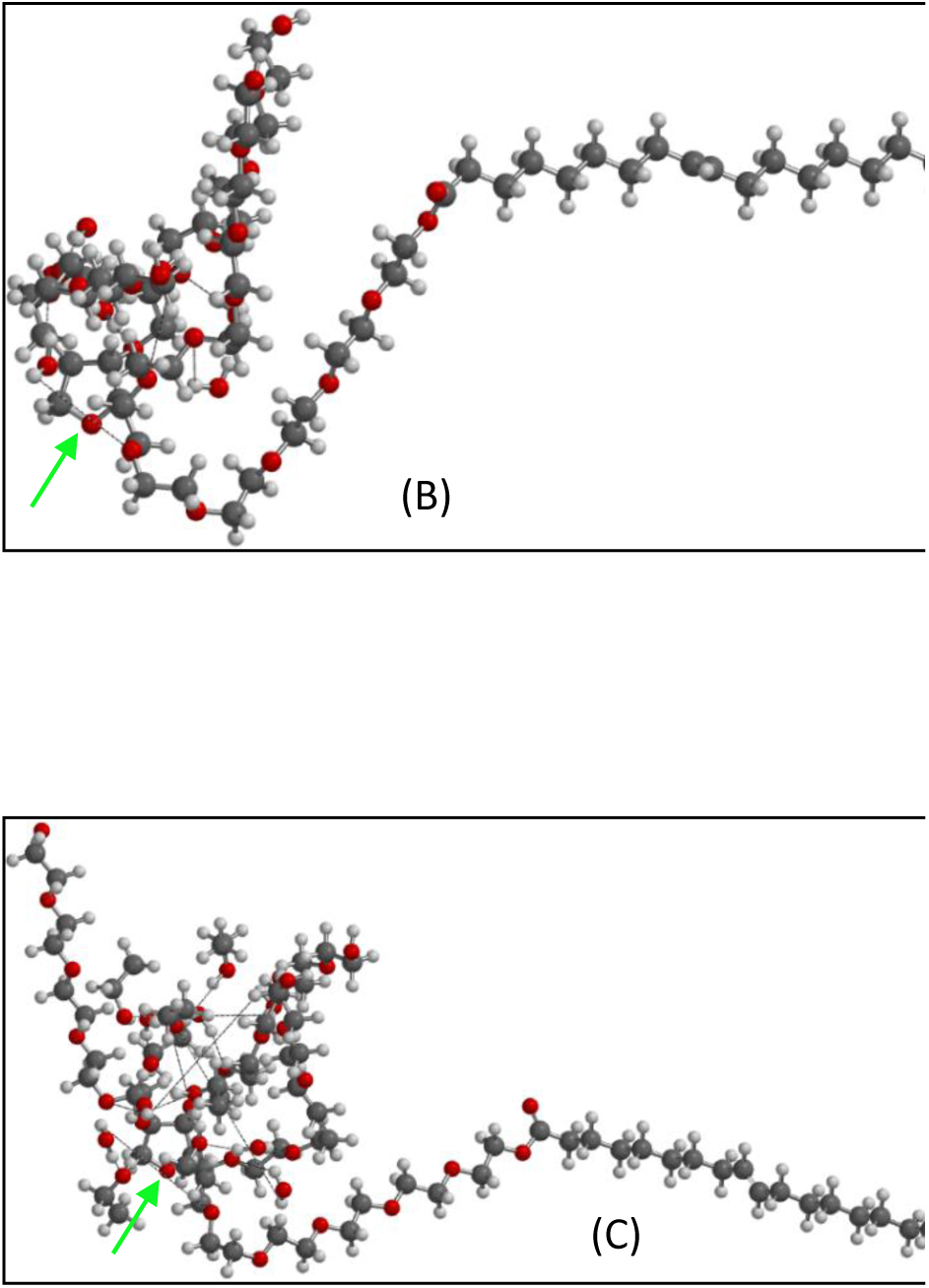
Comparison of (B) P-80 molecule interacting with 7 water molecules at positions 1-7; (C) P-80 molecule interacting with 7 water molecules at positions 1-7; and these in turn interacting with 7 ethanol molecules (CH_3_CH_2_OH).

**Figure 10.**
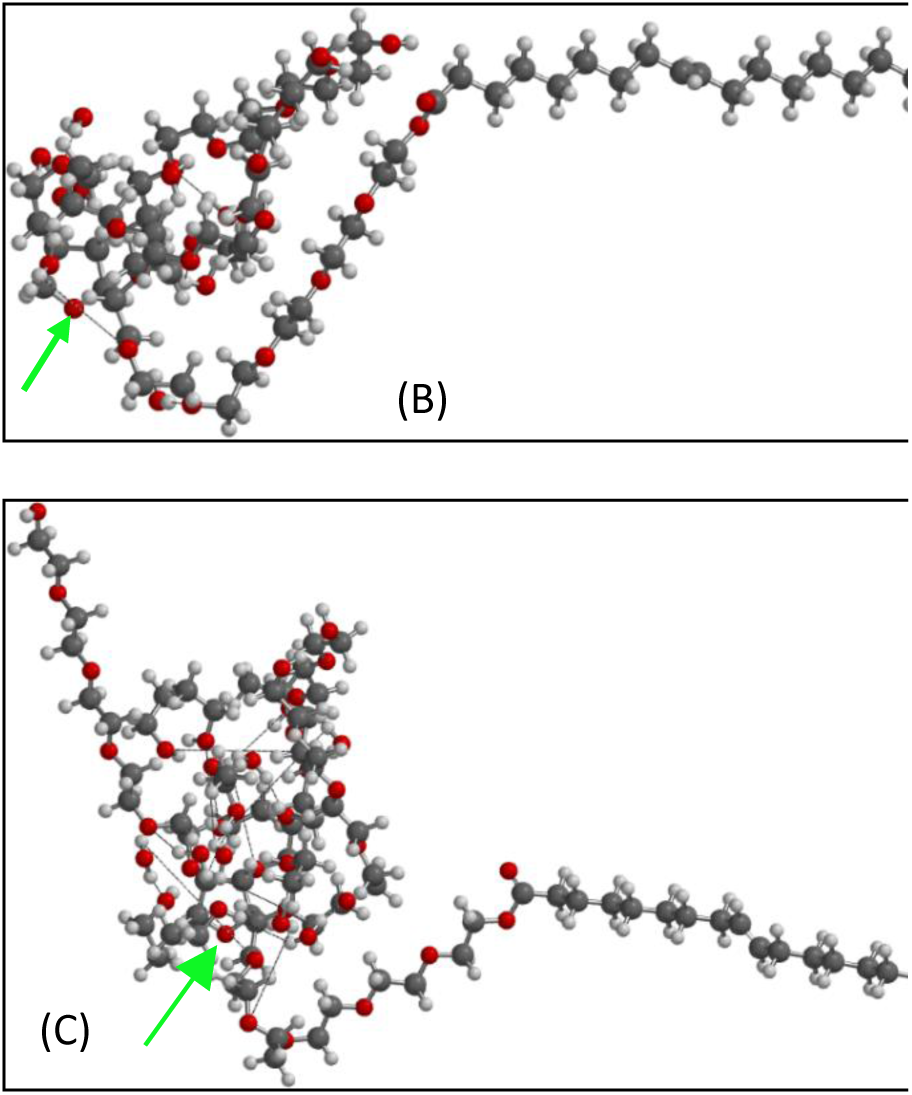
Comparison of (B) P-80 molecule interacting with 8 water molecules at positions 1-8; (C) P-80 molecule interacting with 8 water molecules at positions 1-8; and these in turn interacting with 8 ethanol molecules (CH_3_CH_2_OH).

**Figure 11.**
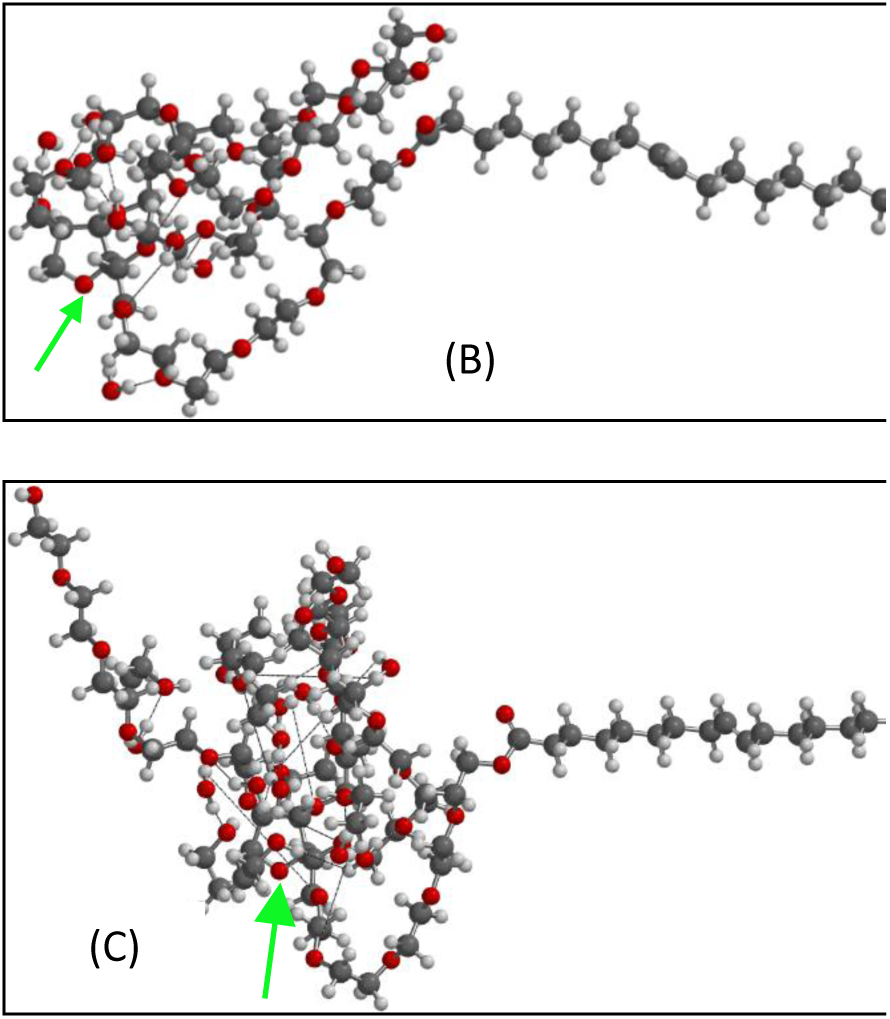
Comparison of (B) P-80 molecule interacting with 9 water molecules at positions 1-9; (C) P-80 molecule interacting with 9 water molecules at positions 1-9; and these in turn interacting with 9 ethanol molecules (CH_3_CH_2_OH).

**Figure 12.**
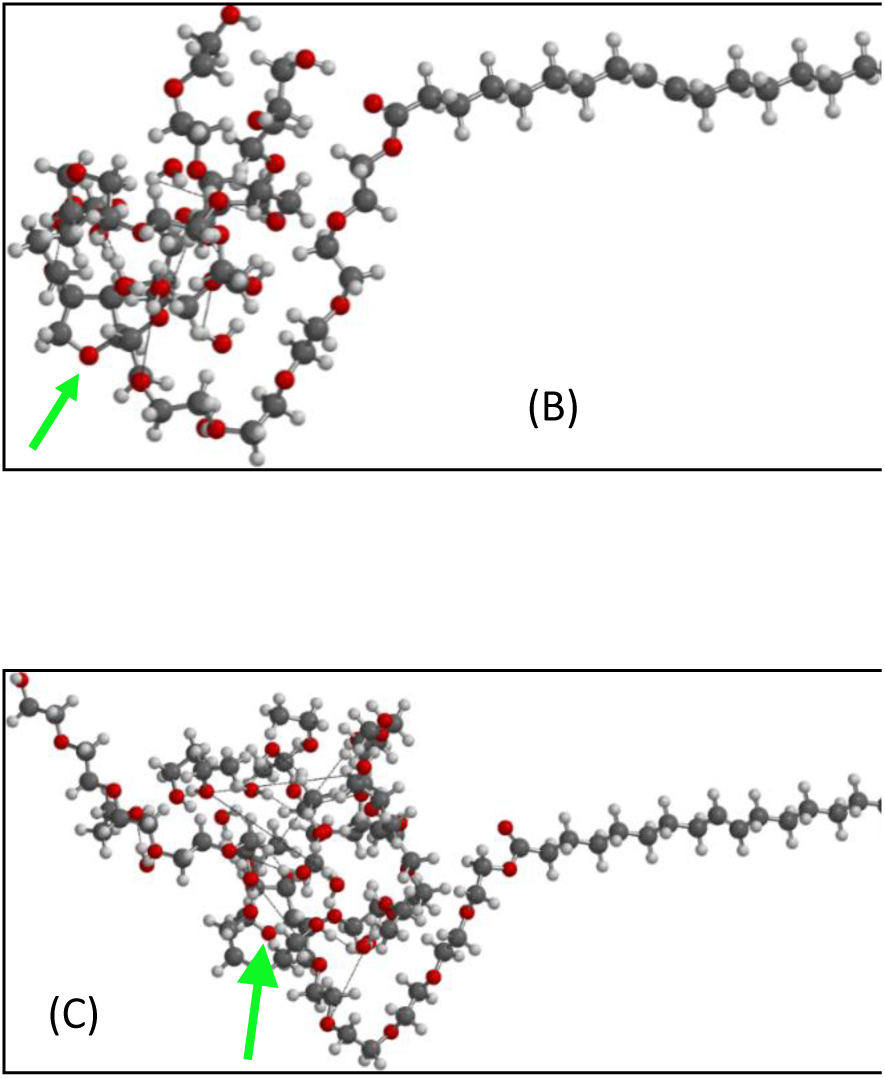
Comparison of: (B) P-80 molecule interacting with 10 water molecules at positions 1-10; (C) P-80 molecule interacting with 10 water molecules at positions 1-10; and these in turn interacting with 10 ethanol molecules (CH_3_CH_2_OH).

**Figure 13.**
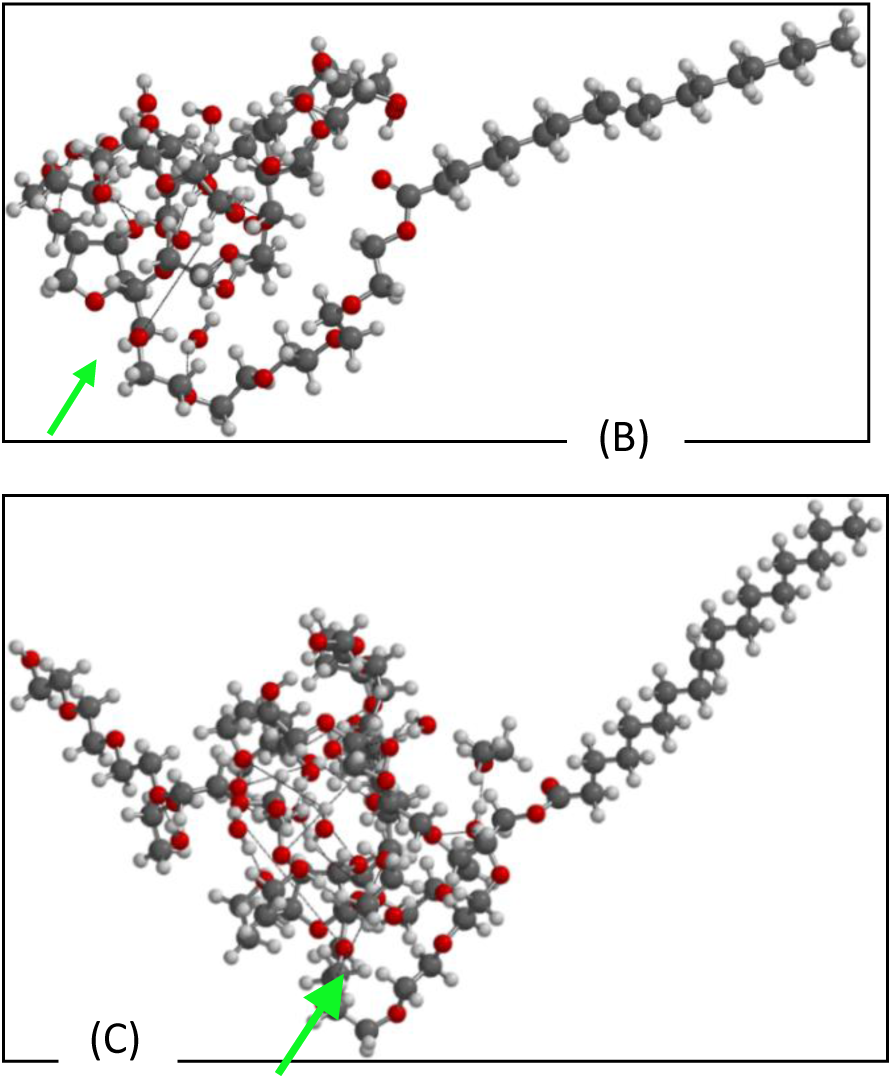
Comparison of (B) P-80 molecule interacting with 11 water molecules at positions 1-11; (C) P-80 molecule interacting with 11 water molecules at positions 1-11; and these in turn interacting with 11 ethanol molecules (CH_3_CH_2_OH).

**Figure 14.**
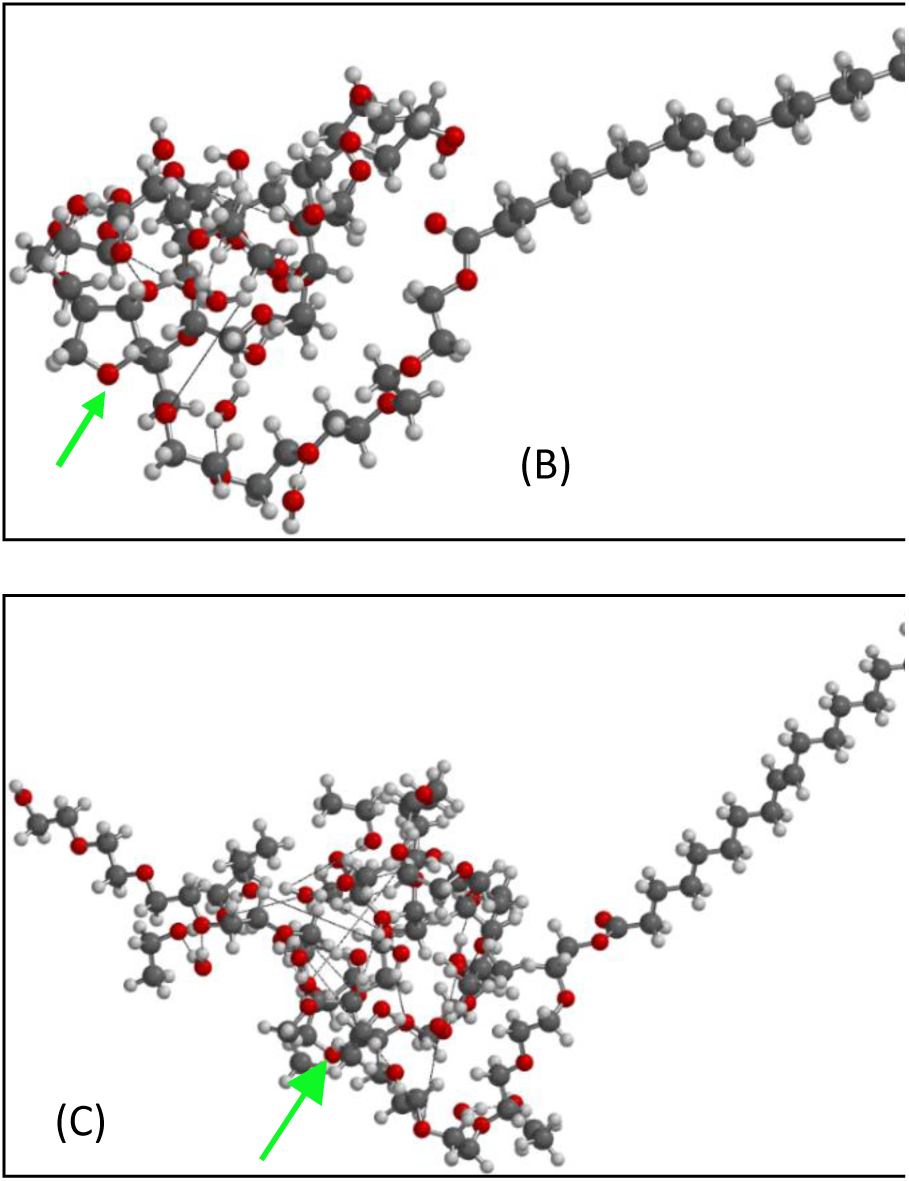
Comparison of (B) P-80 molecule interacting with 12 water molecules at positions 1-12; (C) P-80 molecule interacting with 12 water molecules at positions 1-12; and these in turn interacting with 12 ethanol molecules (CH_3_CH_2_OH).

**Figure 15.**
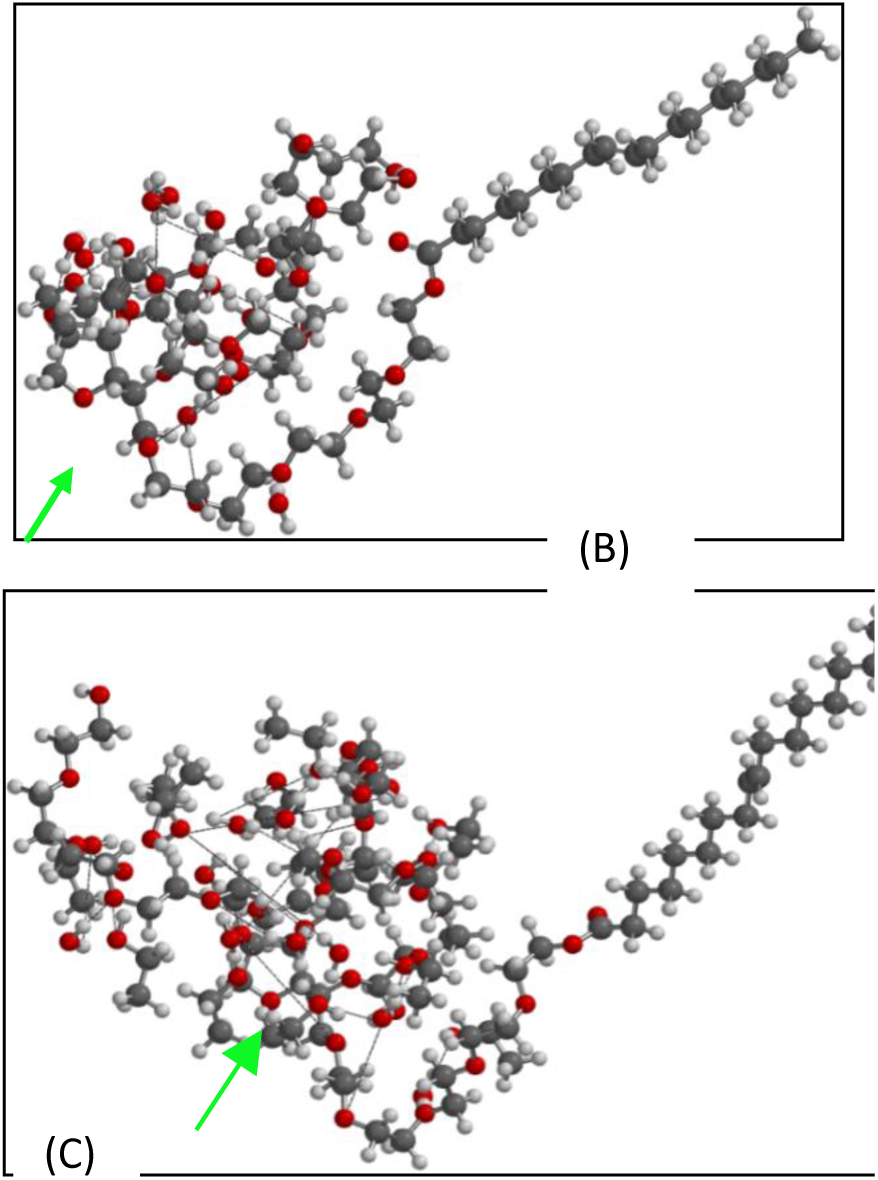
Comparison of (B) P-80 molecule interacting with 13 water molecules at positions 1-13; (C) P-80 molecule interacting with 13 water molecules at positions 1-13; and these in turn interacting with 13 ethanol molecules (CH_3_CH_2_OH).

**Figure 16.**
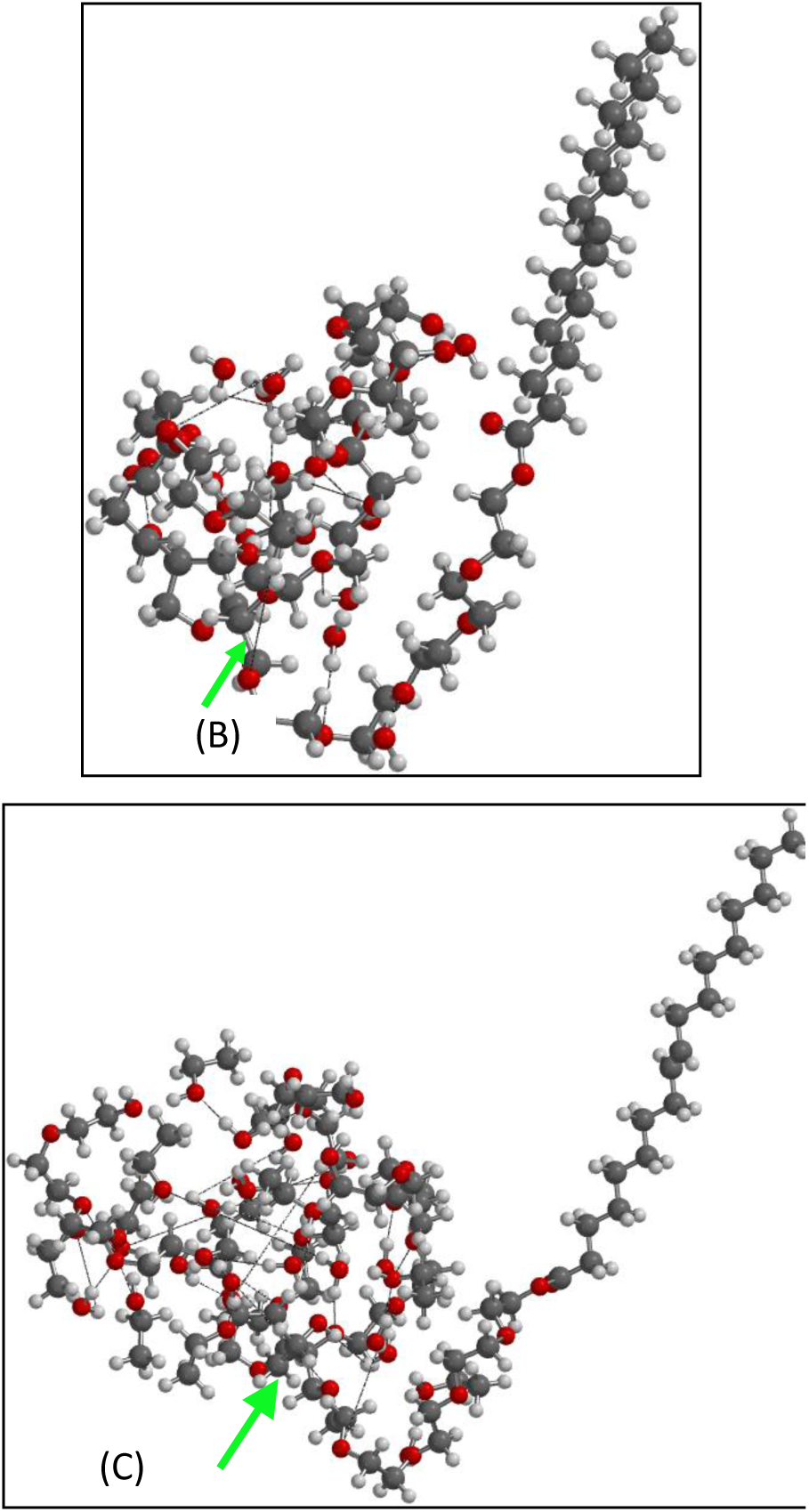
Comparison of (B) P-80 molecule interacting with 14 water molecules at positions 1-14; (C) P-80 molecule interacting with 14 water molecules at positions 1-14; and these in turn interacting with 14 ethanol molecules (CH_3_CH_2_OH).

**Figure 17.**
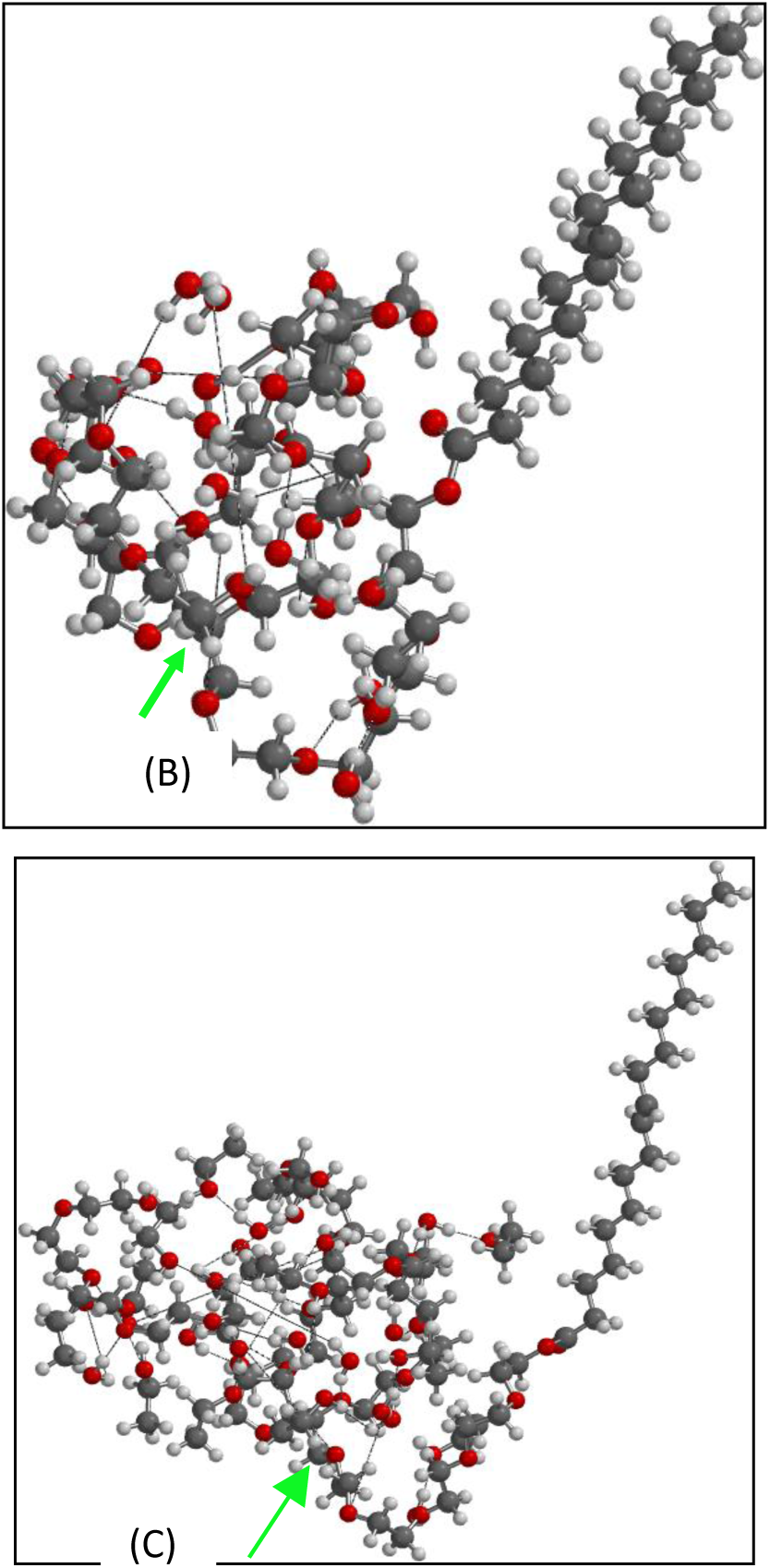
Comparison of (B) P-80 molecule interacting with 15 water molecules at positions 1-15; (C) P-80 molecule interacting with 15 water molecules at positions 1-15; and these in turn interacting with 15 ethanol molecules (CH_3_CH_2_OH).

**Figure 18.**
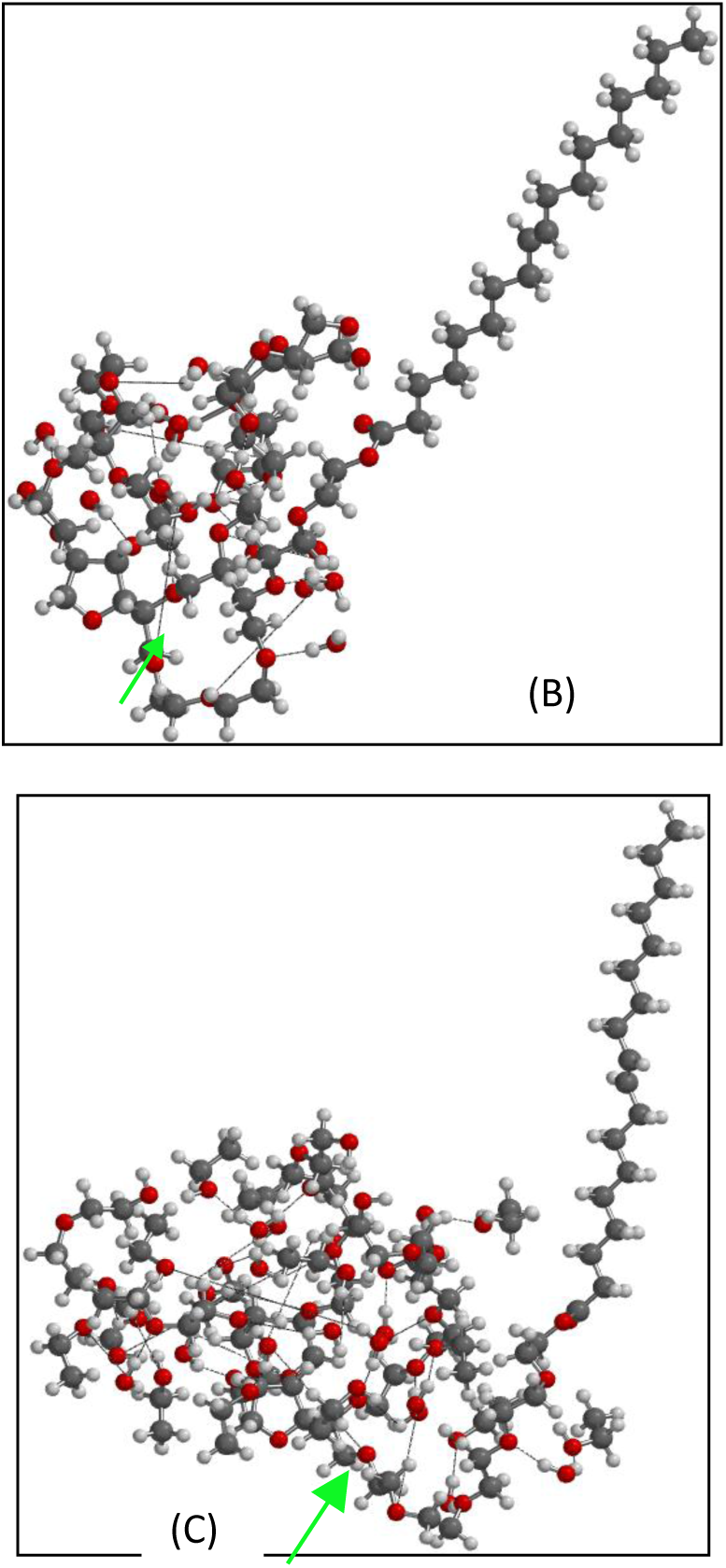
Comparison of (A) P-80 molecule without interaction; (B) P-80 molecule interacting with 16 water molecules at positions 1-16; (C) P-80 molecule interacting with 16 water molecules at positions 1-16; and these in turn interacting with 16 ethanol molecules (CH_3_CH_2_OH).

**Figure 19.**
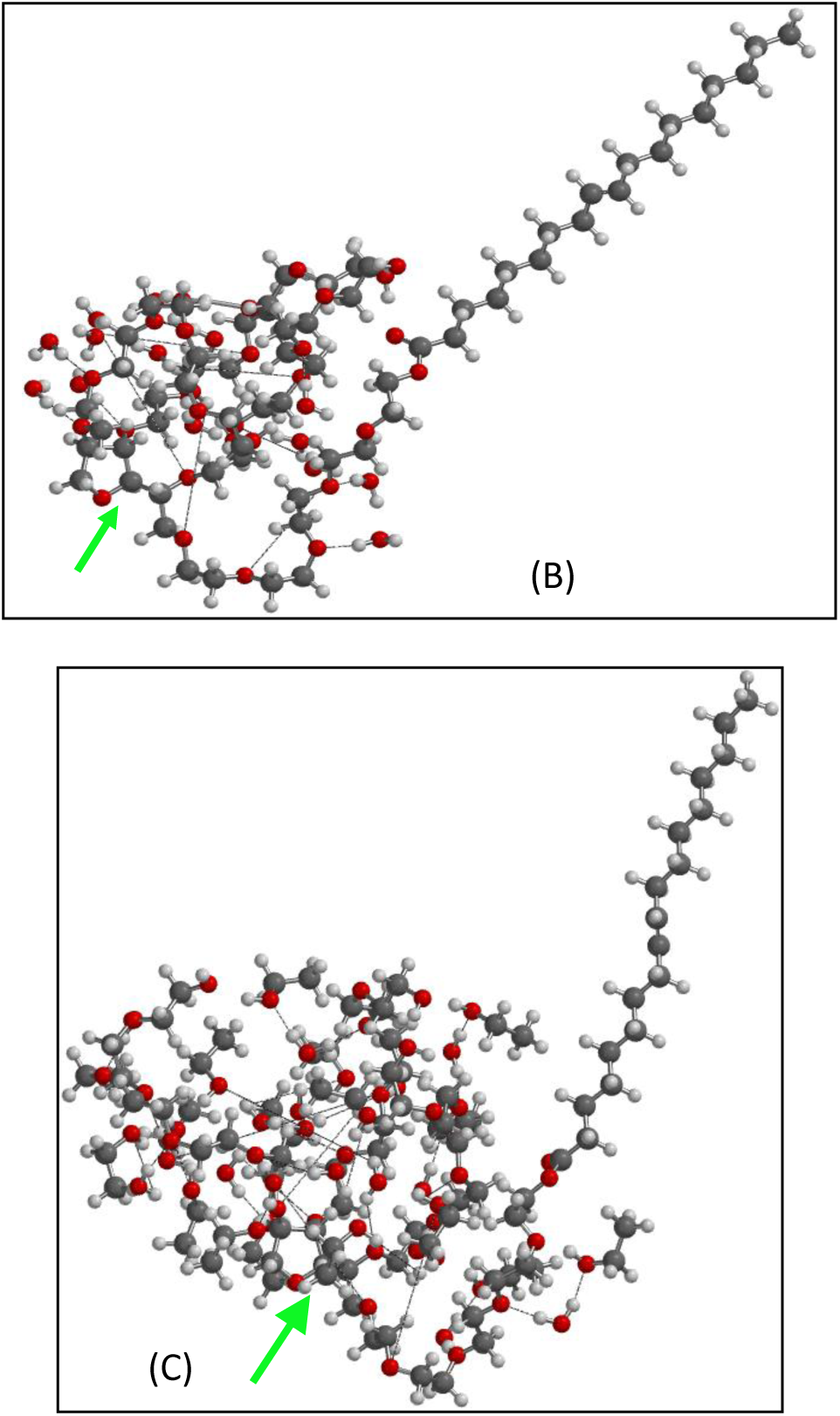
Comparison of (B) P-80 molecule interacting with 17 water molecules at positions 1-17; (C) P-80 molecule interacting with 17 water molecules at positions 1-17; and these in turn interacting with 17 ethanol molecules (CH_3_CH_2_OH).

**Figure 20.**
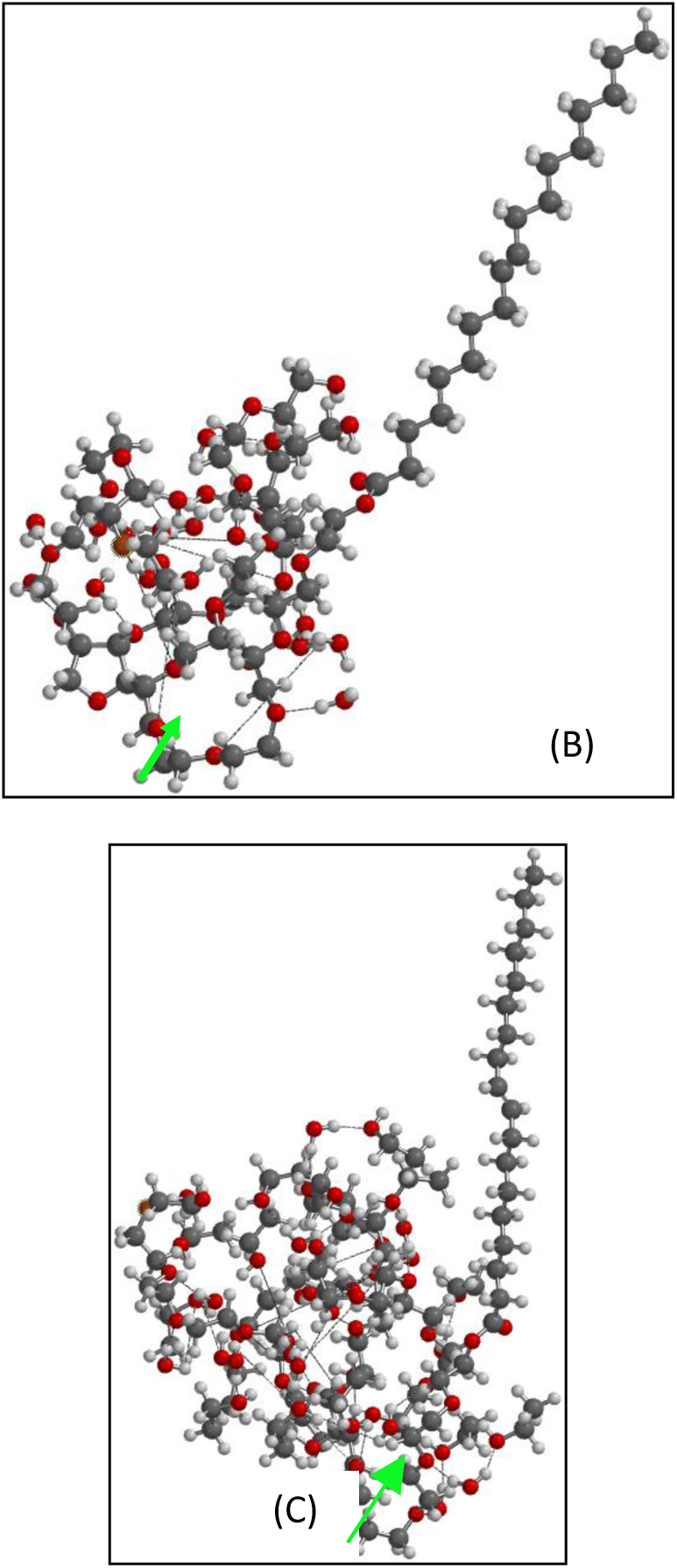
Comparison of (B) P-80 molecule interacting with 18 water molecules at positions 1-18; (C) P-80 molecule interacting with 18 water molecules at positions 1-18; and these in turn interacting with 18 ethanol molecules (CH_3_CH_2_OH).

**Figure 21.**
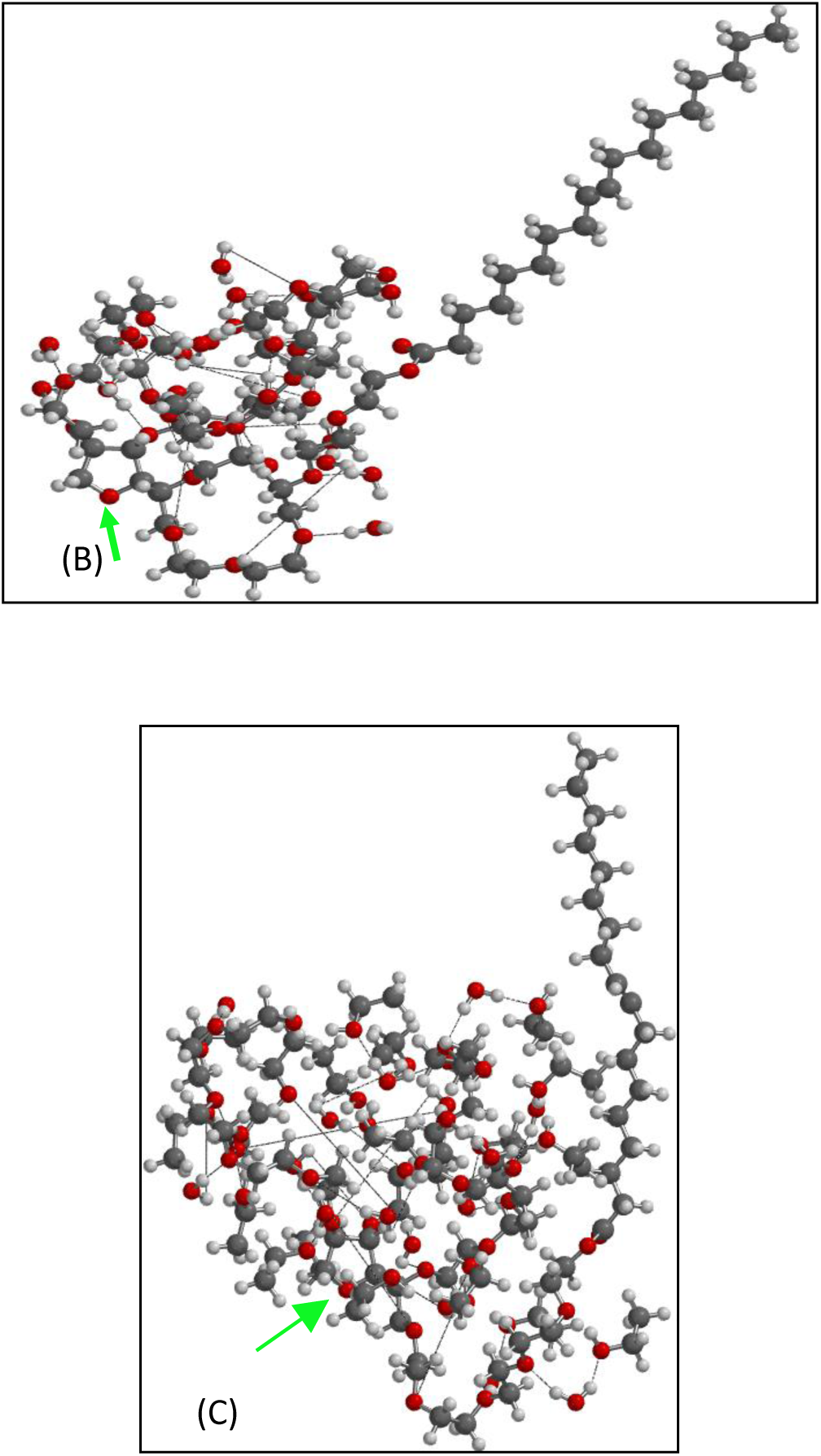
Comparison of (B) P-80 molecule interacting with 19 water molecules at positions 1-19; (C) P-80 molecule interacting with 19 water molecules at positions 1-19; and these in turn interacting with 19 ethanol molecules (CH_3_CH_2_OH).

**Figure 22.**
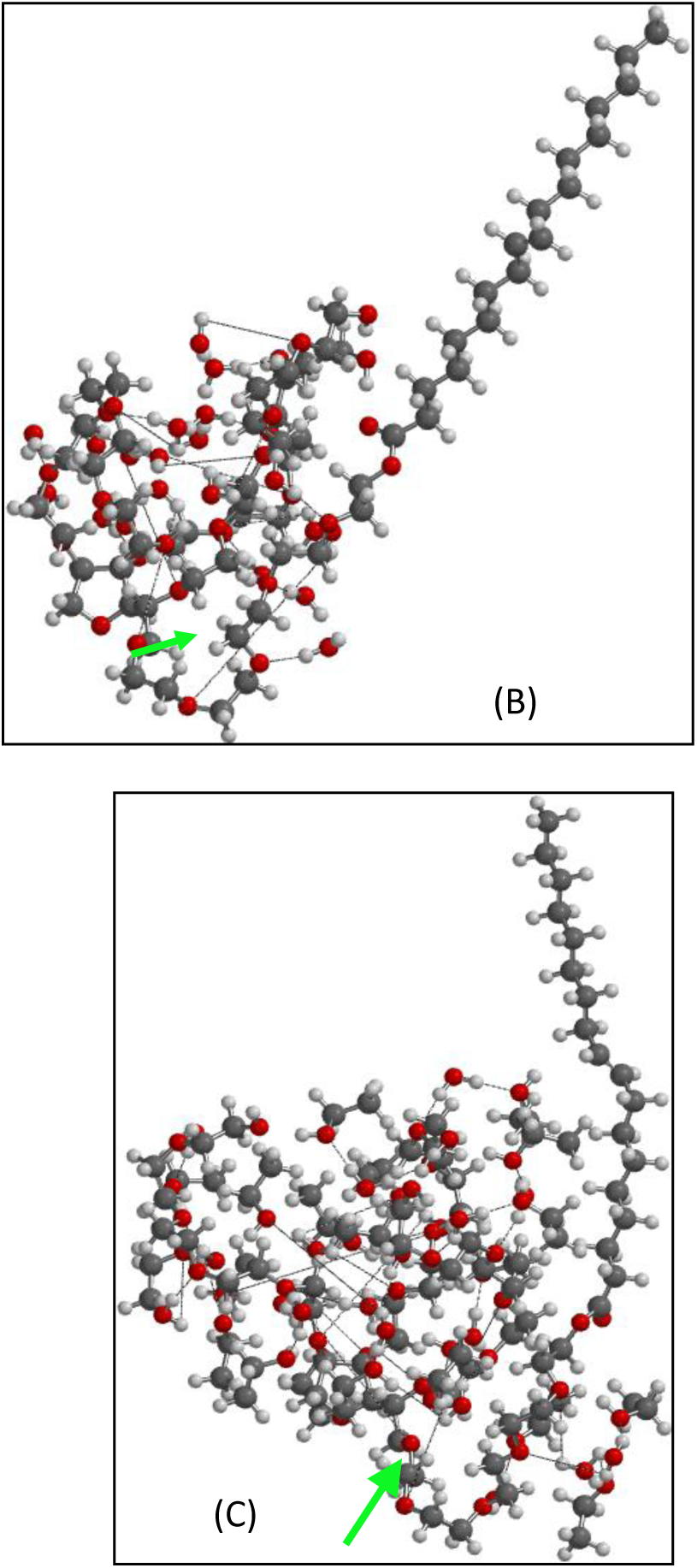
Comparison of (B) P-80 molecule interacting with 20 water molecules at positions 1-20; (C) P-80 molecule interacting with 20 water molecules at positions 1-20; and these in turn interacting with 20 ethanol molecules (CH_3_CH_2_OH).

**Figure 23.**
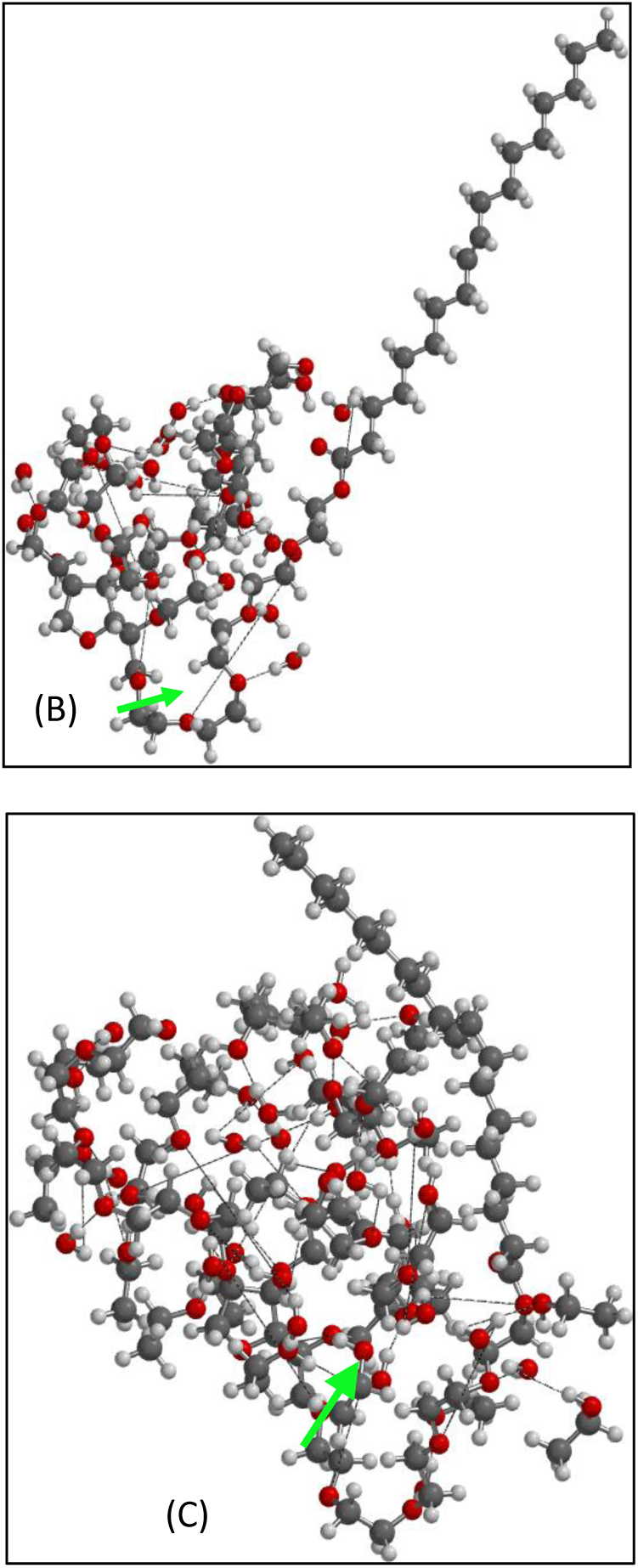
Comparison of (B) P-80 molecule interacting with 22 water molecules at positions 1-21; (C) P-80 molecule interacting with 21 water molecules at positions 1-21; and these in turn interacting with 21 ethanol molecules (CH_3_CH_2_OH).

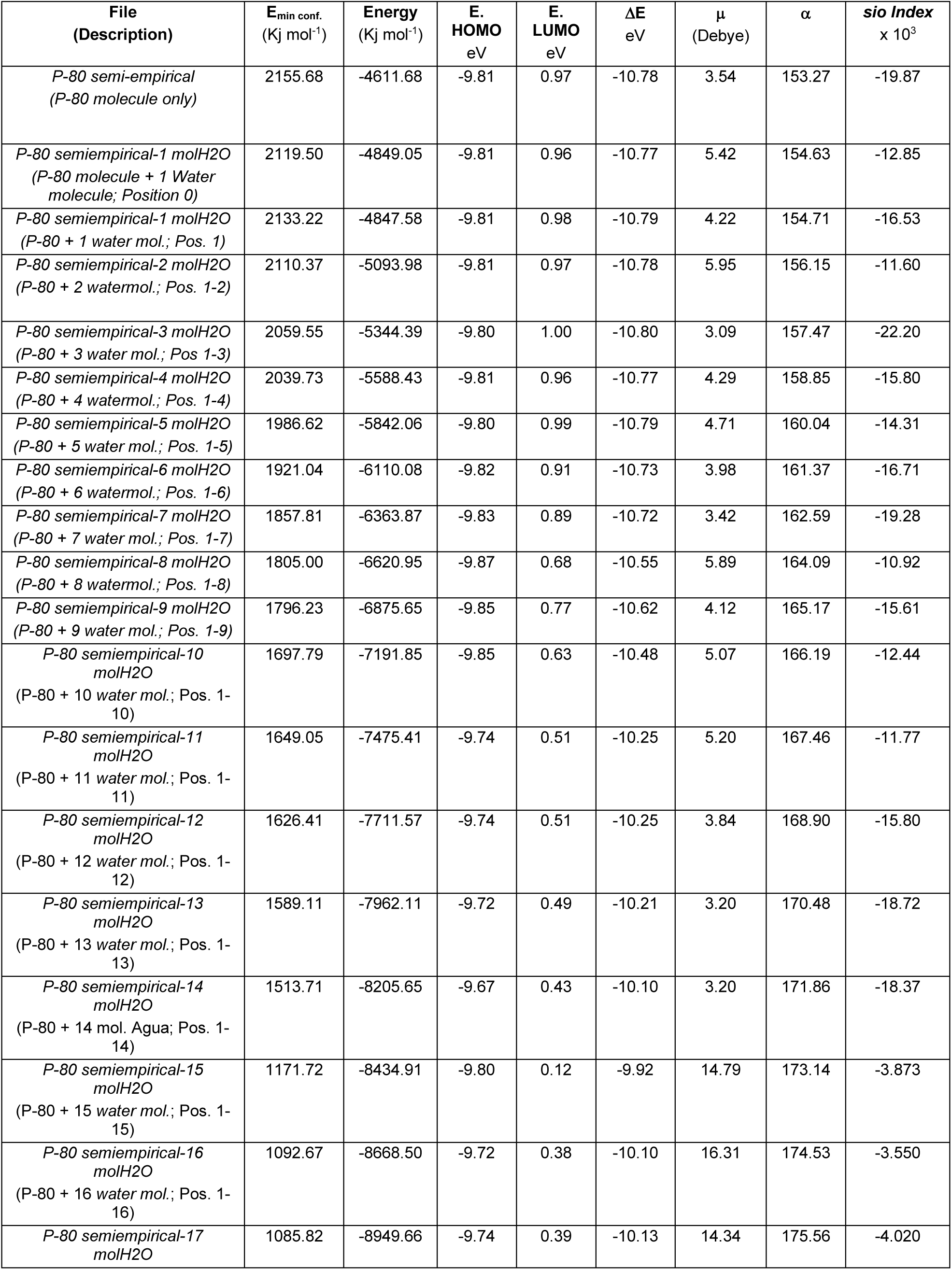
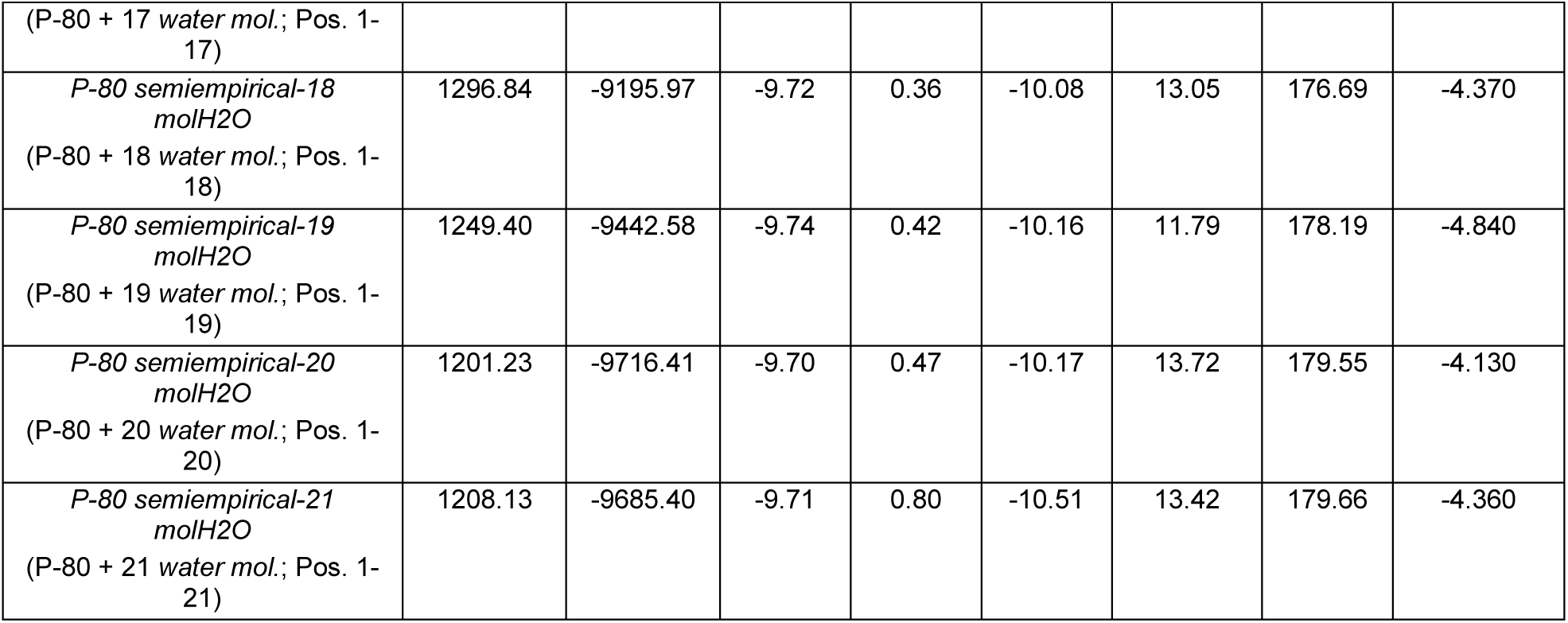
Table 1.

Table 2.

**Table 2**
| **Archive**<br>**(Description)** | **E<sub>min conf.</sub>**<br>**(Kj mol<sup>-1</sup>)** | **Energy**<br>**(Kj mol<sup>-1</sup>)** | **E.**<br>**HOMO**<br>**eV** | **E.**<br>**LUMO**<br>**eV** | **ΔE**<br>**eV** | **μ**<br>**(Debye)** | **α** | **sio**<br>**Index**<br>**x 10<sup>3</sup>** |
| --- | --- | --- | --- | --- | --- | --- | --- | --- |
| *P-80 semi-empirical*<br>*(P-80 molecule alone)* | 2155.68 | -4611.68 | -9.81 | 0.97 | -10.78 | 3.54 | 153.27 | -19.87 |
| *P-80 semiempirical-1 molH<sub>2</sub>O-1mol-Etanol(P-80 + 1 water mol. + 1 ethanol mol.; Pos. 0)* | 2084.25 | -5122.89 | -9.81 | 0.96 | -10.77 | 3.89 | 159.28 | -17.40 |
| *P-80 semiempirical-1 molH<sub>2</sub>O-1mol-Etanol(P-80 + 1 water mol. + 1 ethanol mol.; Pos. 1)* | 2099.22 | -5110.89 | -9.81 | 0.98 | -10.79 | 5.19 | 159.29 | -13.10 |
| *P-80 semiempirical-2 molH<sub>2</sub>O-2 mol-etanol(P-80 + 2 water mol. + 2 ethanol mol.; Pos. 1-2)* | 2017.94 | -5619.04 | -9.80 | 0.99 | -10.79 | 5.35 | 165.33 | -12.20 |
| *P-80 semiempirical-3 molH<sub>2</sub>O-3 mol-etanol(P-80 + 3 water mol. + 3 ethanol mol.; Pos. 1-3)* | 1906.29 | -6153.95 | -9.80 | 1.00 | -10.80 | 5.26 | 170.91 | -12.00 |
| *P-80 semiempirical-4 molH<sub>2</sub>O-4 mol-etanol(P-80 + 4 water mol. + 4 ethanol mol.; Pos. 1-4)* | 1831.20 | -6668.53 | -9.79 | 1.00 | -10.79 | 2.52 | 176.85 | -24.20 |
| *P-80 semiempirical-5 molH<sub>2</sub>O-5 mol-etanol(P-80 + 5 water mol. + 5 ethanol mol.; Pos. 1-5)* | 1693.28 | -7219.60 | -9.81 | 0.95 | -10.76 | 12.17 | 183.01 | -4.800 |
| *P-80 semiempirical-6 molH<sub>2</sub>O-6 mol-etanol(P-80 + 6 water mol. + 6 ethanol mol.; Pos. 1-6)* | 1575.93 | -7732.19 | -9.38 | 0.92 | -10.30 | 7.15 | 188.75 | -7.632 |
| *P-80 semiempirical-7 molH<sub>2</sub>O-7 mol-etanol(P-80 + 7 water mol. + 7 ethanol mol.; Pos. 1-7)* | 1510.67 | -8288.53 | -9.82 | 0.95 | -10.77 | 4.94 | 194.81 | -11.20 |
| *P-80 semiempirical-8 molH<sub>2</sub>O-8 mol-etanol(P-80 + 8 water mol. + 8 ethanol mol.; Pos. 1-8)* | 1394.31 | -8817.74 | -9.83 | 0.85 | -10.68 | 10.05 | 200.48 | -5.300 |
| *P-80 semiempirical-9 molH<sub>2</sub>O-9 mol-etanol(P-80 + 9 water mol. + 9 ethanol mol.; Pos. 1-9)* | 1313.04 | -9332.15 | -9.78 | 1.07 | -10.85 | 6.55 | 206.05 | -8.000 |
| *P-80 semiempirical-10 molH<sub>2</sub>O-10 mol-etanol(P-80 + 10 water mol. + 10 ethanol mol.; Pos. 1-10)* | 1211.90 | -9860.12 | -9.79 | 1.06 | -10.85 | 6.23 | 212.10 | -8.200 |
| *P-80 semiempirical-11 molH<sub>2</sub>O-11 mol-etanol(P-80 + 11 water mol. + 11 ethanol mol.; Pos. 1-11)* | 1099.05 | -10417.25 | -9.78 | 1.02 | -10.80 | 5.01 | 38.83 | -55.50 |
| *P-80 semiempirical-12 molH<sub>2</sub>O-12 mol-etanol(P-80 + 12 water mol. + 12 ethanol mol.; Pos. 1-12)* | 1050.08 | -10913.72 | -9.77 | 1.05 | -10.82 | 6.81 | 38.83 | -40.90 |
| *P-80 semiempirical-13 molH<sub>2</sub>O-13 mol-etanol(P-80 + 13 water mol. + 13 ethanol mol.; Pos. 1-13)* | 930.43 | -11438.12 | -9.78 | 0.98 | -10.76 | 4.15 | 38.84 | -66.80 |
| *P-80 semiempirical-14 molH<sub>2</sub>O-14 mol-etanol(P-80 + 14 water mol. + 14 ethanol mol.; Pos. 1-14)* | 836.34 | -12015.56 | -9.77 | 0.99 | -10.76 | 8.99 | 38.84 | -30.80 |
| *P-80 semiempirical-15 molH<sub>2</sub>O-15 mol-etanol(P-80 + 15 water mol. + 15 ethanol mol.; Pos. 1-15)* | 773.31 | -12504.63 | -9.86 | 0.86 | -10.72 | 5.47 | 38.85 | -50.40 |
| *P-80 semiempirical-16 molH<sub>2</sub>O-16 mol-etanol(P-80 + 16 water mol. + 16 ethanol mol.; Pos. 1-16)* | 695.57 | -12996.73 | -9.88 | 0.73 | -10.61 | 9.13 | 38.88 | -29.90 |
| *P-80 semiempirical-17 molH<sub>2</sub>O-17 mol-etanol(P-80 + 17 water mol. + 17 ethanol mol.; Pos. 1-17)* | 579.66 | -13523.93 | -9.77 | 0.95 | -10.72 | 8.68 | 38.85 | -31.80 |
| *P-80 semiempirical-18 molH<sub>2</sub>O-18 mol-etanol(P-80 + 18 water mol. + 18 ethanol mol.; Pos. 1-18)* | 512.75 | -14053.51 | -9.86 | 0.83 | -10.69 | 9.45 | 38.86 | -29.10 |
| *P-80 semiempirical-19 molH<sub>2</sub>O-19 mol-etanol(P-80 + 19 water mol. + 19 ethanol mol.; Pos. 1-19)* | 411.46 | -14577.21 | -10.10 | 0.68 | -10.78 | 8.48 | 38.84 | -32.70 |
| *P-80 semiempirical-20 molH<sub>2</sub>O-20 mol-etanol(P-80 + 20 water mol. + 20 ethanol mol.; Pos. 1-20)* | 341.52 | -15097.11 | -10.09 | 0.67 | -10.76 | 11.86 | 38.84 | -23.40 |
| *P-80 semiempirical-21 molH<sub>2</sub>O-21 mol-etanol(P-80 + 21 water mol. + 21 ethanol mol.; Pos. 1-21)* | 220.70 | -15519.30 | -7.51 | 0.93 | -8.44 | 23.64 | 39.38 | -9.10 |

### Table 3, Figures 2 (D) – 23 (D) (The arrow indicates the position of sorbitol)

In the case of Series 3, in which Polysorbate 80 is occupied with 22 water molecules and ethanol molecules are added, the value of P is very high up to 6 ethanol molecules. In this case the same reasoning applies, but with a high concentration of water and a low amount of ions up to 6 ethanols and a little higher from 7 ethanols onwards with improved density.

The results of the three Series seem to demonstrate that the proportion of Water and ions in relation to the other components of the Biocompatible Mixtures is important.

The criterion issued for all molecular forms with DPM below the P-80 value (4.06), but with a P value above 39, is valid in the same way as saying that they would be candidates to influence the improvement and recovery of crude oil under conditions of very low ionic concentration, which could be achieved by increasing the amount of water, sacrificing the improvement of Viscosity and improving Density or vice versa. This vice versa would define which molecular forms improve Viscosity and which improve Density, namely: those that are more likely to be found within the crude oil at low ratios of water and ions with respect to other components, would improve Viscosity. 2. – And vice versa for Density.

What is clear is that it is not the absolute quantities or concentrations, but, for example, the ratios: Water: P-80; P-80: Water; Water: Ethanol, the ionic concentration, etc.

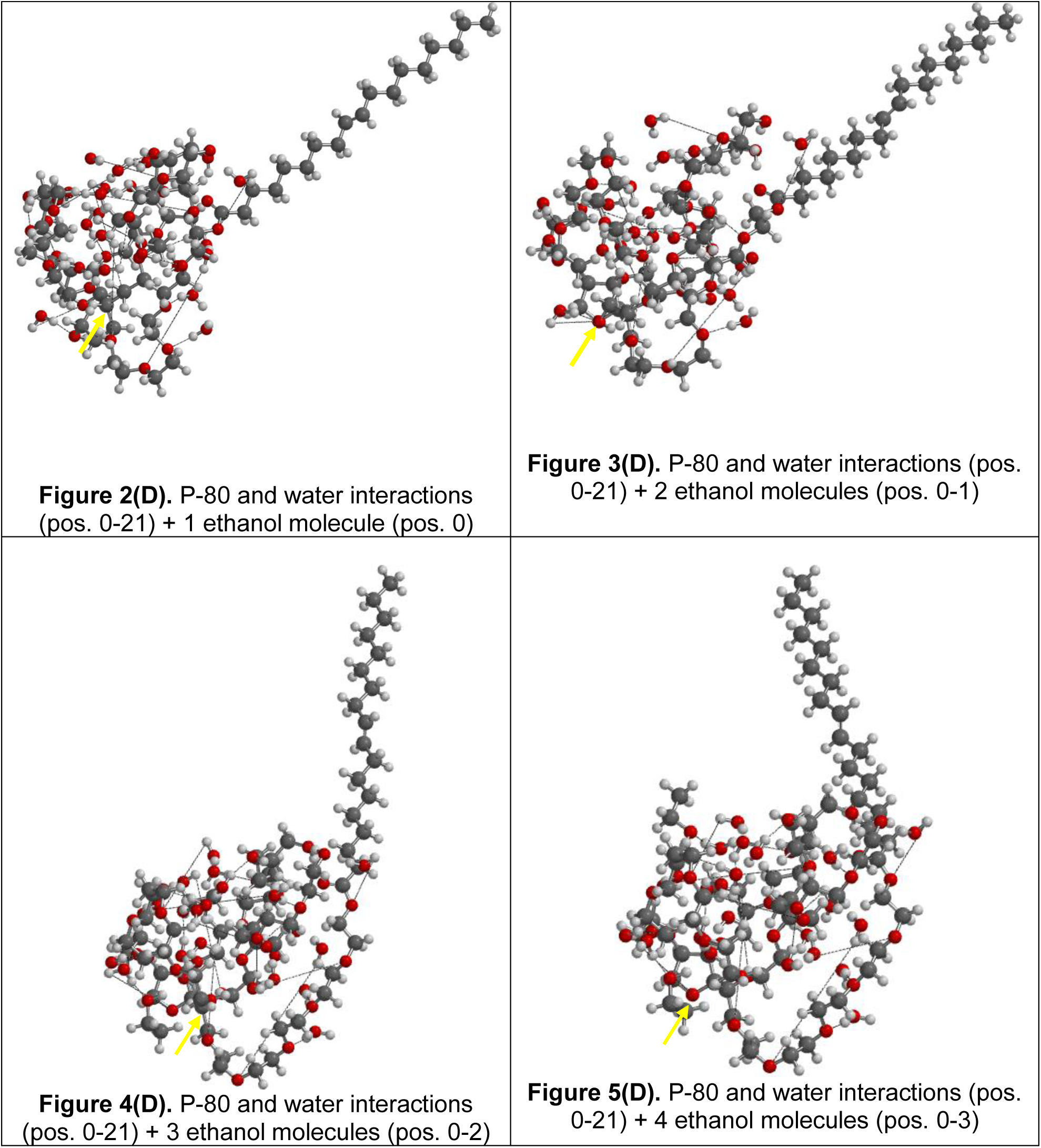
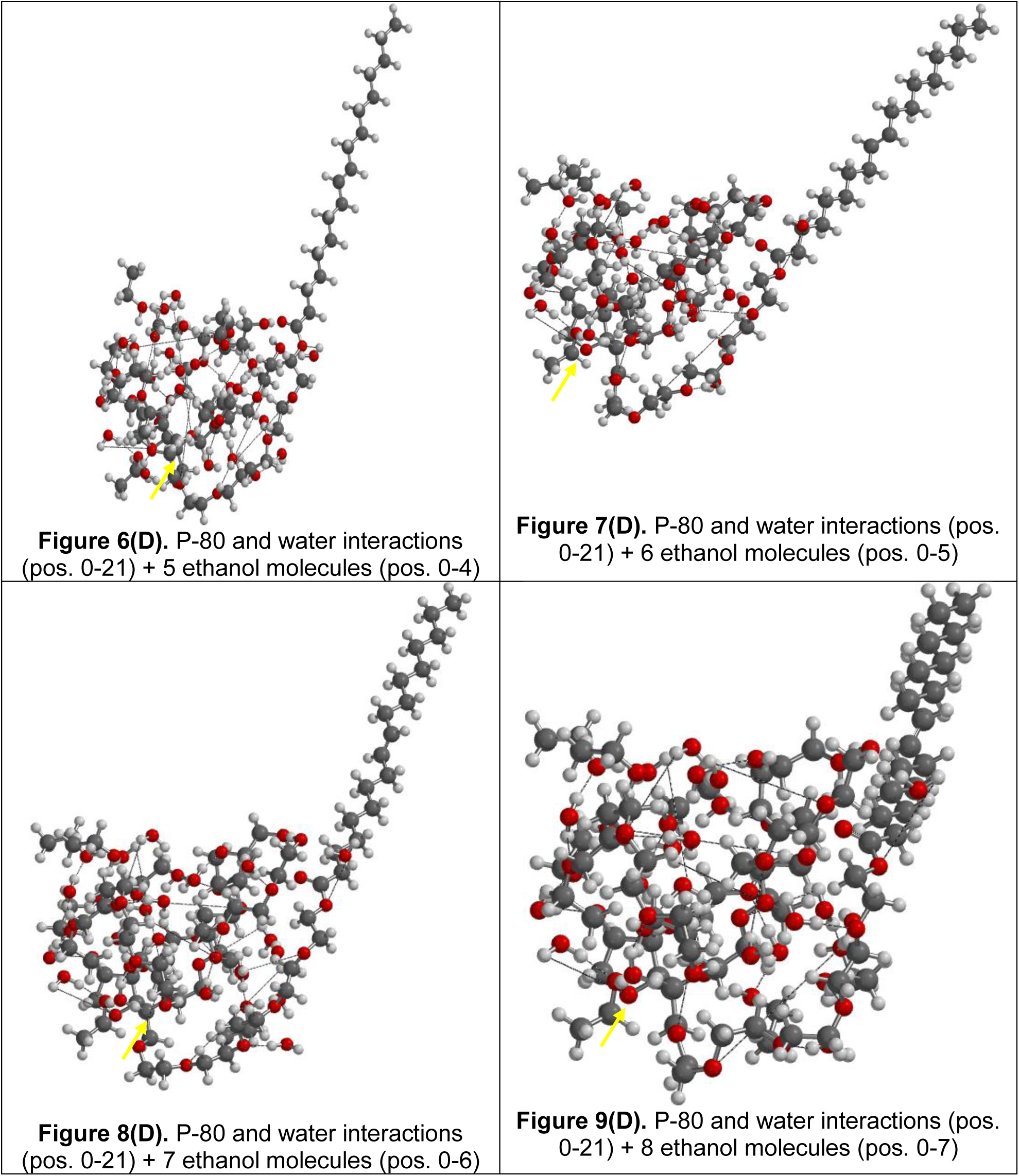
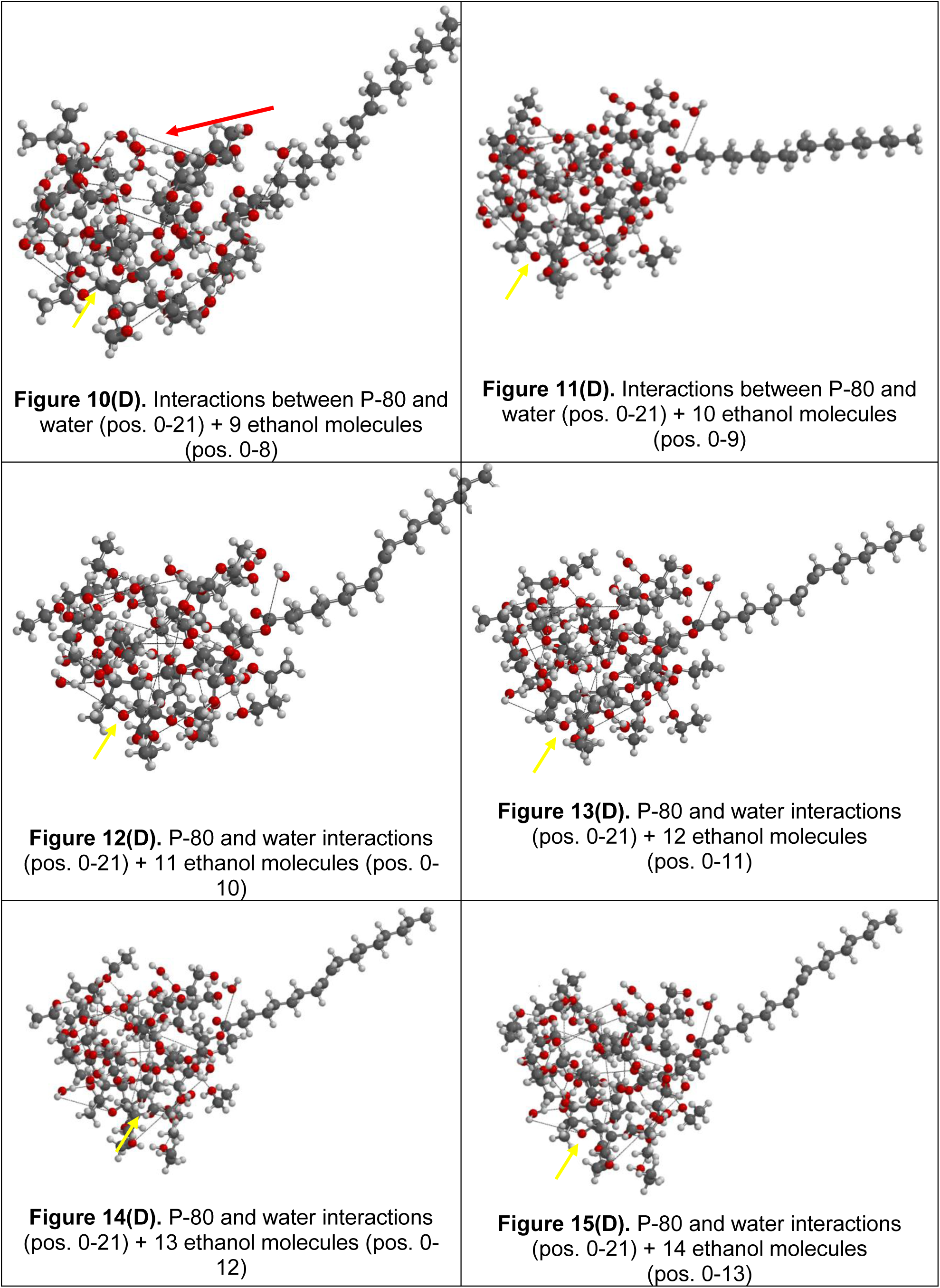
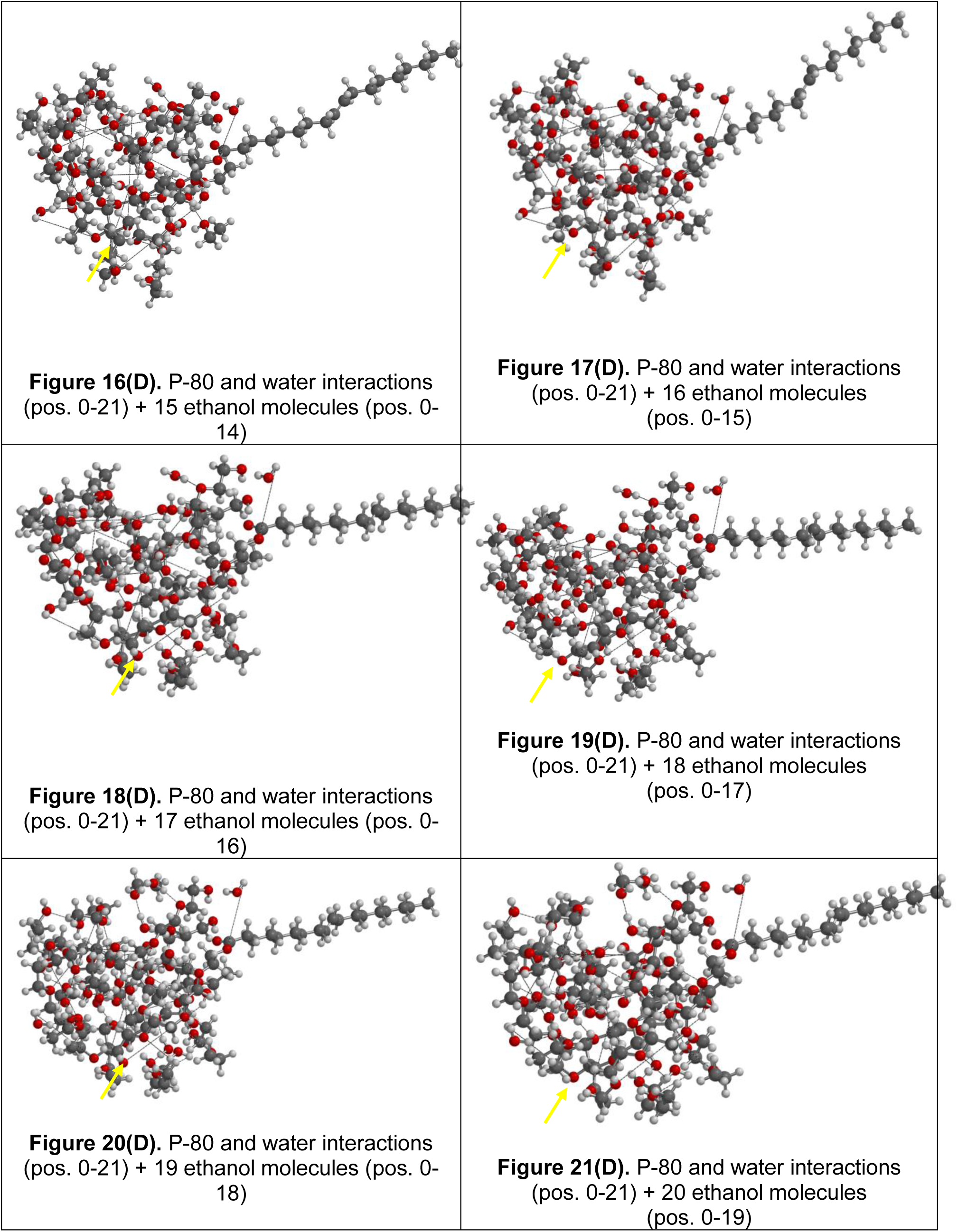
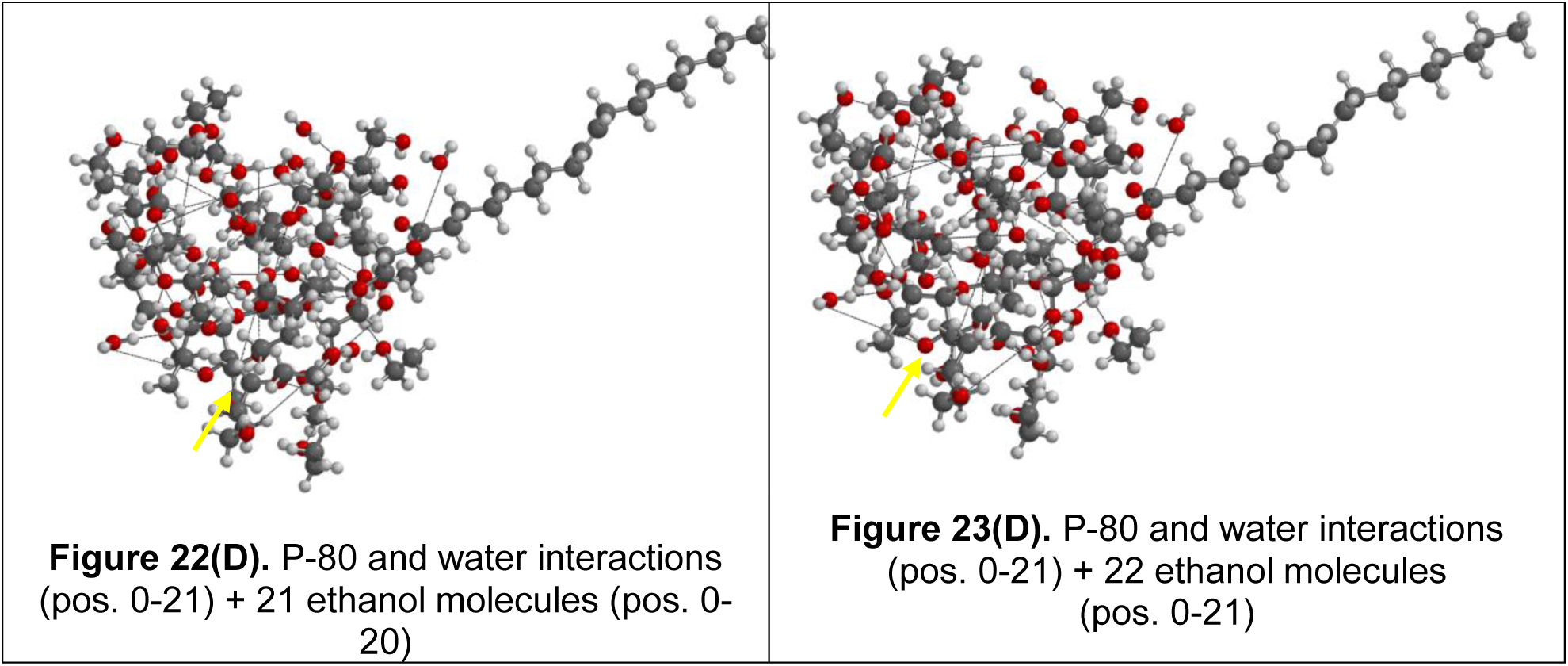

Table 3.

**Table 3**
| **Archive**<br>**(Description)** | **E<sub>min conf.</sub>**<br>**(Kj mol<sup>-1</sup>)** | **Energy**<br>**(Kj mol<sup>-1</sup>)** | **E. HOMO**<br>**eV** | **E. LUMO**<br>**eV** | **ΔE**<br>**eV** | **μ**<br>**(Debye)** | **α** | **sio**<br>**Index**<br>**x 10<sup>3</sup>** |
| --- | --- | --- | --- | --- | --- | --- | --- | --- |
| *P-80 semiempirical*<br>*(molécula de P-80 solo)* | 2155.68 | -4611.68 | -9.81 | 0.97 | -10.78 | 3.54 | 153.27 | -19.87 |
| (0) +P-80 semiempirical-21<br>molH<sub>2</sub>O [P-80 + 22 Water<br>molecules. at positions 1-21] | 1208.13 | -9685.40 | -9.71 | 0.80 | -10.51 | 13.42 | 179.66 | -4.360 |
| (0) +P-80 semiempirical-21<br>molH<sub>2</sub>O+ (0) ethanolmol.<br>[P-80 + 22 Watermol., Pos. 0-<br>21 + 1 ethanolmol., Pos. 0] | 1054.50 | -10475.57 | -9.76 | 0.70 | -10.46 | 12.75 | 186.76 | -4.390 |
| (0-1) +P-80 semiempirical-21<br>molH<sub>2</sub>O+ (0-1) ethanolmol.<br>[P-80 + 22 water mol., Pos. 0-<br>21 + 2 Ethanol mol., Pos. 0-1] | 1024.40 | -10751.00 | -9.76 | 0.64 | -10.40 | 12.00 | 191.65 | -4.522 |
| (0-2) +P-80 semiempirical-21<br>molH<sub>2</sub>O+ (0-2) ethanolmol<br>[P-80 + 22 water mol., Pos. 0-<br>21 + 3 ethanol mol., Pos. 0-2] | 984.24 | -10992.73 | -9.84 | 0.62 | -10.46 | 4.72 | 196.45 | -11.28 |
| (0-3) +P-80 semiempirical-21<br>molH<sub>2</sub>O+ (0-3) ethanol mol<br>[P-80 + 22 water mol., Pos. 0-<br>21 + 4 ethanol mol., Pos. 0-3] | 928.05 | -11270.86 | -9.80 | 0.67 | -10.47 | 4.29 | 200.80 | -12.15 |
| (0-4) +P-80 semiempirical-21<br>molH<sub>2</sub>O+ (0-4) ethanol mol<br>[P-80 + 22 water mol., Pos. 0-<br>21 + 5 ethanol mol., Pos. 0-4] | 893.28 | -11528.41 | -9.79 | 0.66 | -10.45 | 6.31 | 205.21 | -8.070 |
| (0-5) +P-80 semiempirical-21<br>molH<sub>2</sub>O+ (0-5) ethanol mol<br>[P-80 + 22 water mol., Pos. 0-<br>21 + 6 ethanol mol., Pos. 0-5] | 859.22 | -11807.24 | -9.79 | 0.78 | -10.57 | 2.90 | 209.71 | -17.38 |
| (0-6) +P-80 semiempirical-21<br>molH<sub>2</sub>O+ (0-6) ethanol mol<br>[P-80 + 22 water mol., Pos. 0-<br>21 + 7 Ethanol mol., Pos. 0-6] | 795.68 | -12066.93 | -9.79 | 0.65 | -10.44 | 9.23 | 38.91 | -29.06 |
| (0-7) +P-80 semiempirical-21<br>molH<sub>2</sub>O+ (0-7) ethanol mol<br>[P-80 + 22 water mol., Pos. 0-<br>21 + 8 Ethanol mol., Pos. 0-7] | 686.03 | -12378.01 | -9.86 | 0.58 | -10.44 | 6.42 | 38.92 | -41.78 |
| (0-8) +P-80 semiempirical-21<br>molH<sub>2</sub>O+ (0-8) ethanol mol<br>[P-80 + 22 water mol., Pos. 0- | 664.92 | -12646.03 | -9.80 | 0.87 | -10.67 | 3.09 | 38.86 | -88.85 |
| 21 + 9 ethanol mol., Pos. 0-8] |  |  |  |  |  |  |  |  |
| (0-9) +P-80 semiempirical-21 molH<sub>2</sub>O+ (0-9) ethanol mol<br>[P-80 + 22 water mol., Pos. 0-21 + 10 ethanol mol., Pos. 0-9] | 535.75 | -12860.63 | -9.86 | 0.51 | -10.37 | 7.25 | 38.93 | -36.74 |
| (0-10) +P-80 semiempirical-21 molH<sub>2</sub>O+ (0-10) ethanol mol<br>[P-80 + 22 water mol., Pos. 0-21 + 11 ethanol mol., Pos. 0-10] | 518.58 | -13134.73 | -9.81 | 0.70 | -10.51 | 4.91 | 38.90 | -55.02 |
| (0-11) +P-80 semiempirical-21 molH<sub>2</sub>O+ (0-11) ethanol mol.<br>[P-80 + 22 water mol., Pos. 0-21 + 12 ethanol mol., Pos. 0-11] | 395.62 | -13387.03 | -9.83 | 0.65 | -10.48 | 8.98 | 38.91 | -29.99 |
| (0-12) +P-80 semiempirical-21 molH<sub>2</sub>O+ (0-12) ethanol mol.<br>[P-80 + 22 water mol., Pos. 0-21 + 13 ethanol mol., Pos. 0-12] | 355.45 | -13664.91 | -9.79 | 0.75 | -10.54 | 10.56 | 38.89 | -25.66 |
| (0-13) +P-80 semiempirical-21 molH<sub>2</sub>O+ (0-13) ethanol mol.<br>[P-80 + 22 water mol., Pos. 0-21 + 14 ethanol mol., Pos. 0-13] | 335.99 | -13930.30 | -9.78 | 0.76 | -10.54 | 7.80 | 38.89 | -34.74 |
| (0-14) +P-80 semiempirical-21 molH<sub>2</sub>O+ (0-14) ethanol mol.<br>[P-80 + 22 water mol., Pos. 0-21 + 15 ethanol mol., Pos. 0-14] | 298.38 | -14195.92 | -9.79 | 0.80 | -10.59 | 8.46 | 38.88 | -32.19 |
| (0-15) +P-80 semiempirical-21 molH<sub>2</sub>O+ (0-15) ethanol mol.<br>[P-80 + 22 water mol., Pos. 0-21 + 16 ethanol mol., Pos. 0-15] | 259.64 | -14462.59 | -9.78 | 0.79 | -10.57 | 9.41 | 38.89 | -28.88 |
| (0-16) +P-80 semiempirical-21 molH<sub>2</sub>O+ (0-16) ethanol mol.<br>[P-80 + 22 water mol., Pos. 0-21 + 17 ethanol mol., Pos. 0-16] | 205.28 | -14747.78 | -9.82 | 0.65 | -10.47 | 7.40 | 38.91 | -36.36 |
| (0-17) + P-80 semiempirical-21 molH<sub>2</sub>O+ (0-17) ethanol mol.<br>[P-80 + 22 mol. water, Pos. 0-21 + 18 ethanol mol., Pos. 0-17] | 160.14 | -15013.56 | -9.82 | 0.65 | -10.47 | 9.17 | 38.91 | -29.34 |
| (0-18) + P-80 semiempirical-21 molH<sub>2</sub>O+ (0-18) ethanol mol.<br>[P-80 + 22 water mol., Pos. 0-21 + 19 ethanol mol., Pos. 0-18] | 110.19 | -15282.51 | -9.81 | 0.67 | -10.48 | 10.36 | 38.91 | -25.99 |
| (0-19) + P-80 semiempirical-21 molH<sub>2</sub>O+ (0-19) ethanol mol.<br>[P-80 + 22 water mol., Pos. 0-21 + 20 ethanol mol., Pos. 0-19] | 110.14 | -15301.23 | -9.82 | 0.63 | -10.45 | 9.67 | 38.91 | -27.77 |
| (0-20) + P-80 semiempirical- | 259.69 | -14462.60 | -9.78 | 0.79 | -10.57 | 9.36 | 38.89 | -29.03 |
| 21 molH<sub>2</sub>O+ (0-20) ethanol mol.<br>[P-80 + 22 water mol., Pos. 0-21 + 21 ethanol mol., Pos. 0-20] |  |  |  |  |  |  |  |  |
| (0-21) + P-80 semiempirical-21 molH<sub>2</sub>O+ (0-21) ethanol mol.<br>[P-80 + 22 water mol., Pos. 0-21 + 22 ethanol mol., Pos. 0-21] | 259.69 | -14462.59 | -9.78 | 0.79 | -10.57 | 9.38 | 38.89 | -28.97 |

**Figure 24.**
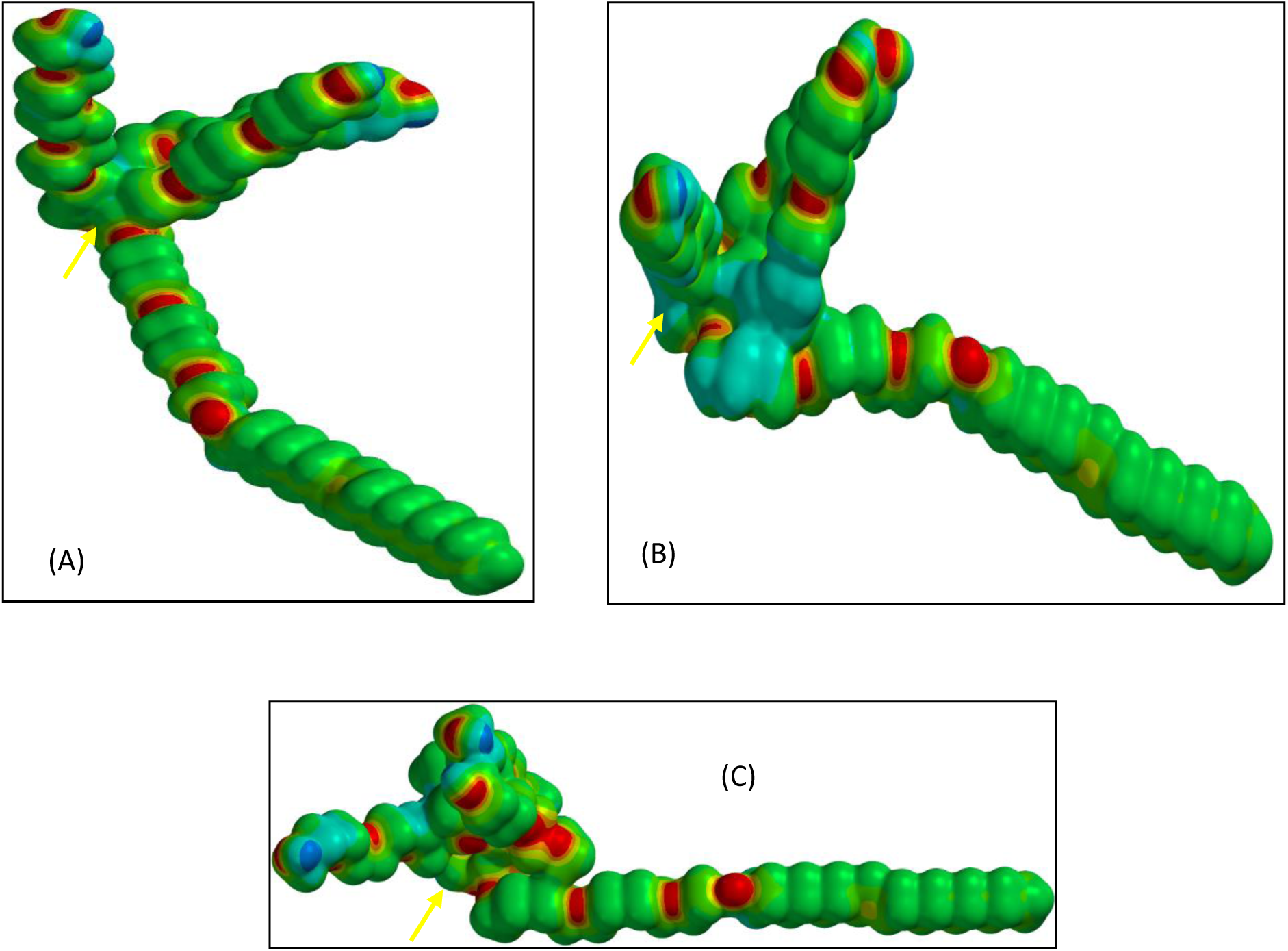
Comparison of Electrostatic Potential Maps (EPM) (A) P-80 molecule without interaction; (B) P-80 molecule interacting with 3 water molecules at positions 1-3; (C) P-80 molecule interacting with 3 water molecules at positions 1-3; and this in turn interacting with 3 ethanol molecules (CH_3_CH_2_OH) at the same sites.

**Figure 25.**
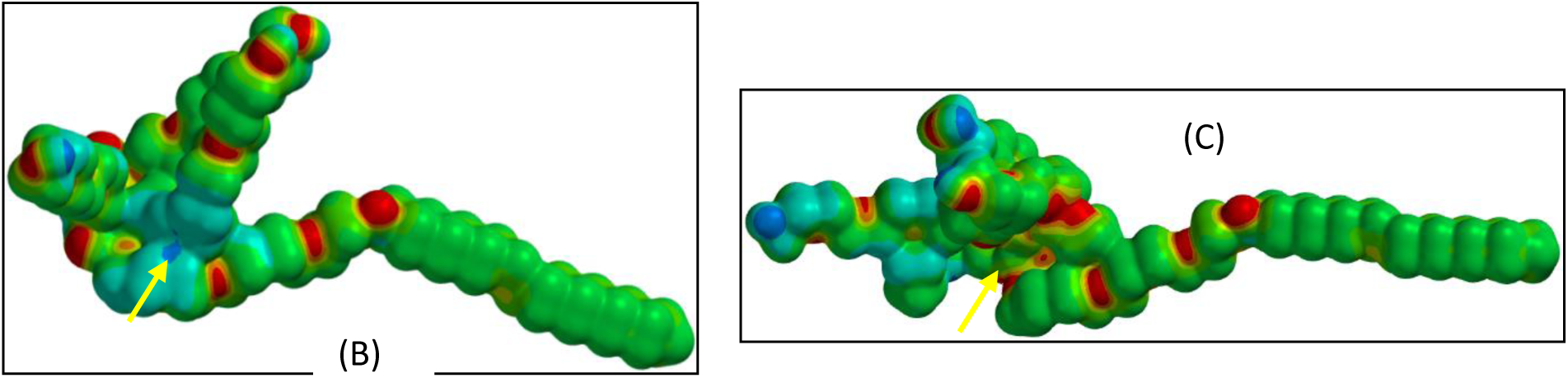
Comparison of Electrostatic Potential Maps (EPM) (B) P-80 molecule interacting with 4 water molecules at positions 1-4; (C) P-80 molecule interacting with 4 water molecules at positions 1-4; and this in turn interacting with 4 ethanol molecules (CH_3_CH_2_OH) at the same locations.

**Figure 26.**
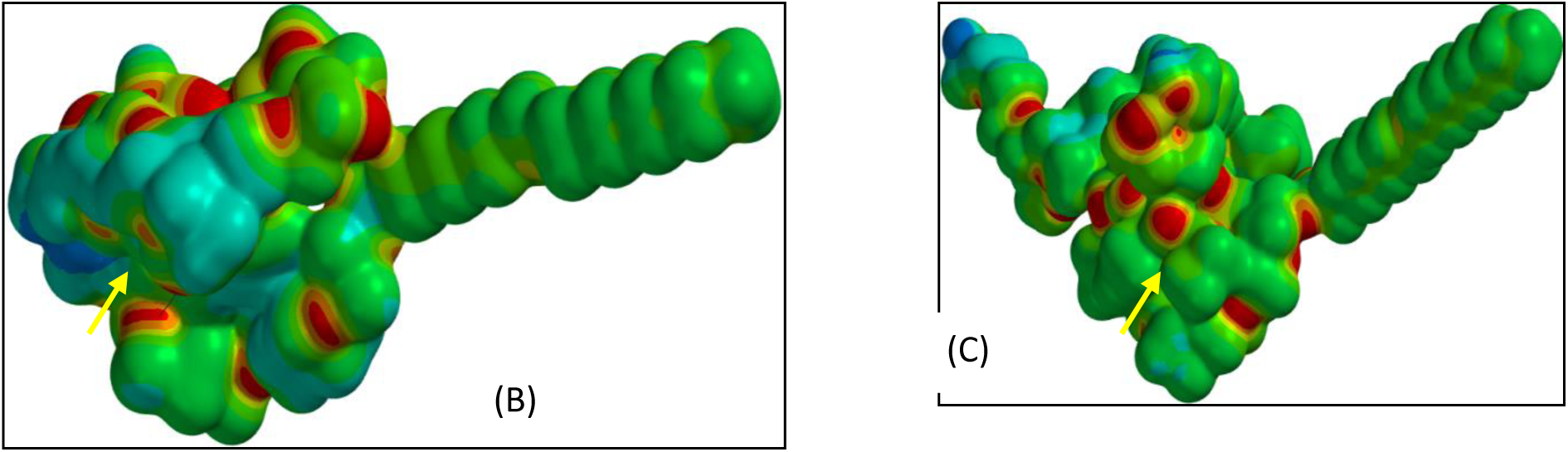
Comparison of Electrostatic Potential Maps (EPM) (B) P-80 molecule interacting with 11 water molecules at positions 1-11; (C) P-80 molecule interacting with 11 water molecules at positions 1-11; and this in turn interacting with 11 ethanol (CH_3_CH_2_OH) molecules at the same locations.

**Figure 27.**
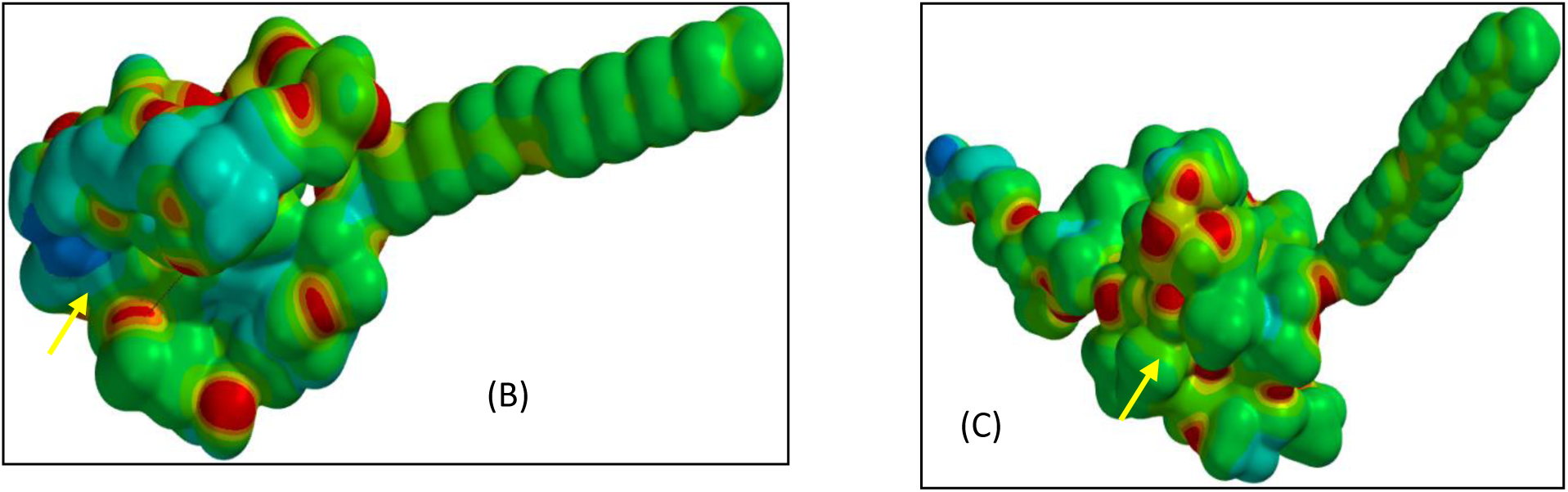
Comparison of Electrostatic Potential Maps (EPM) (B) P-80 molecule interacting with 12 water molecules at positions 1-12; (C) P-80 molecule interacting with 12 water molecules at positions 1-12; and this in turn interacting with 12 ethanol (CH_3_CH_2_OH) molecules at the same locations.

**Figure 28.**
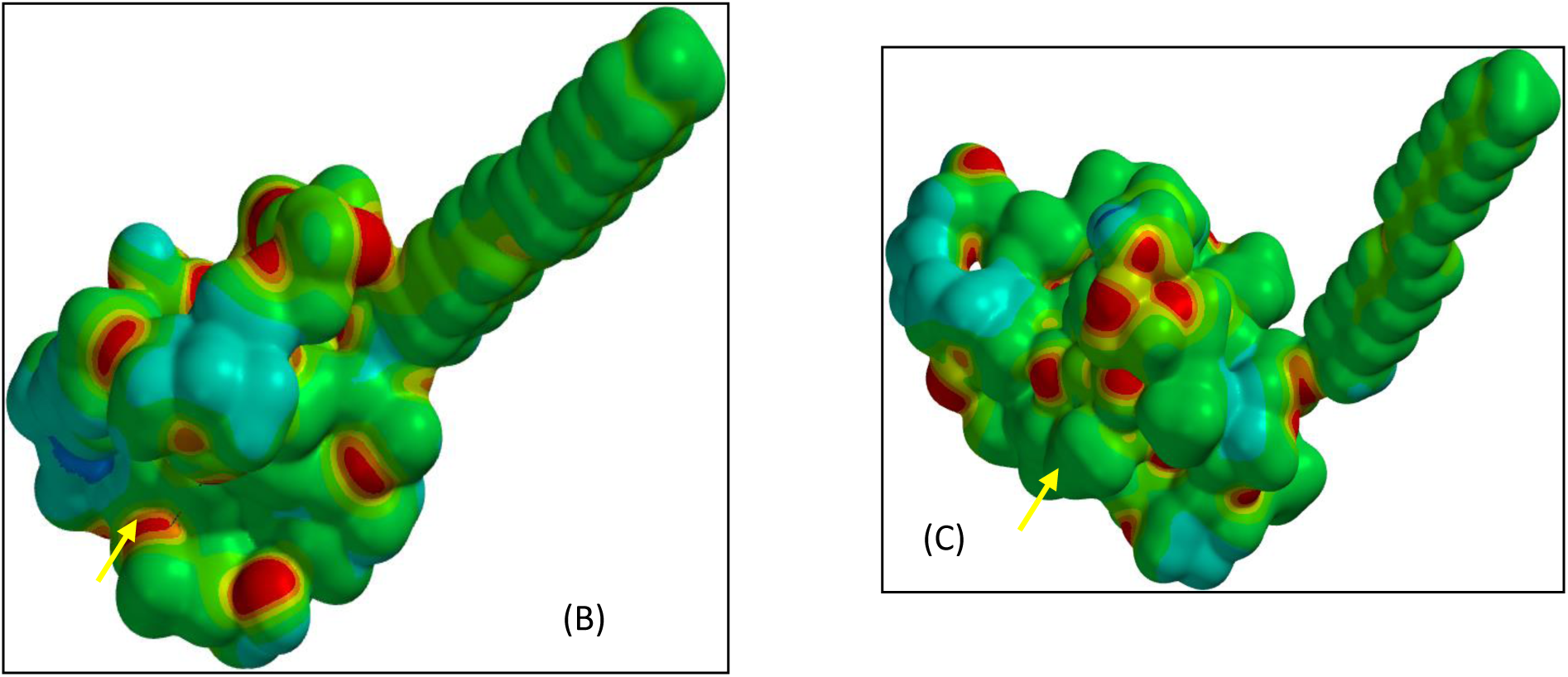
Comparison of Electrostatic Potential Maps (EPM) (B) P-80 molecule interacting with 13 water molecules at positions 1-13; (C) P-80 molecule interacting with 13 water molecules at positions 1-13; and this in turn interacting with 13 ethanol (CH_3_CH_2_OH) molecules at the same locations.

**Figure 29.**
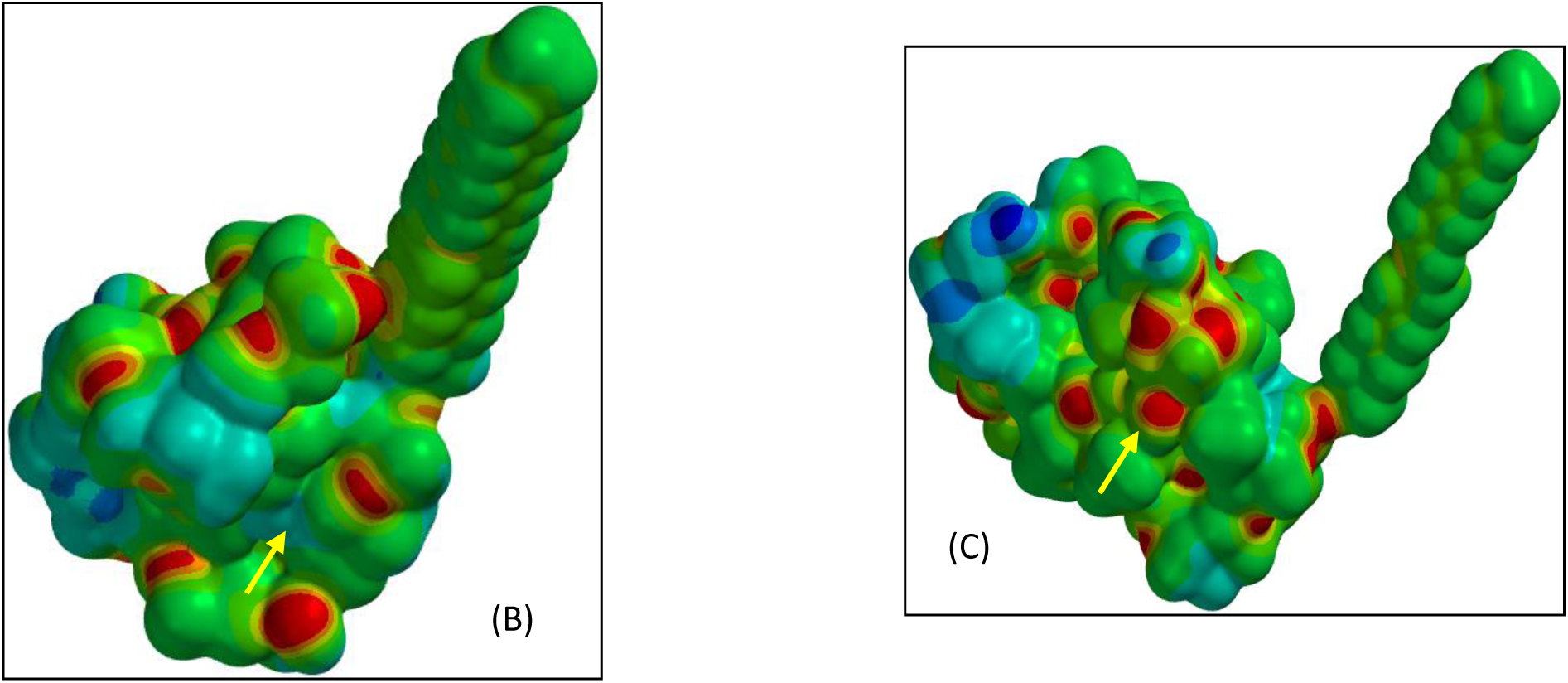
Comparison of Electrostatic Potential Maps (EPM) (B) P-80 molecule interacting with 14 water molecules at positions 1-14; (C) P-80 molecule interacting with 14 water molecules at positions 1-14; and this in turn interacting with 14 ethanol (CH_3_CH_2_OH) molecules at the same locations.

**Figure 30.**
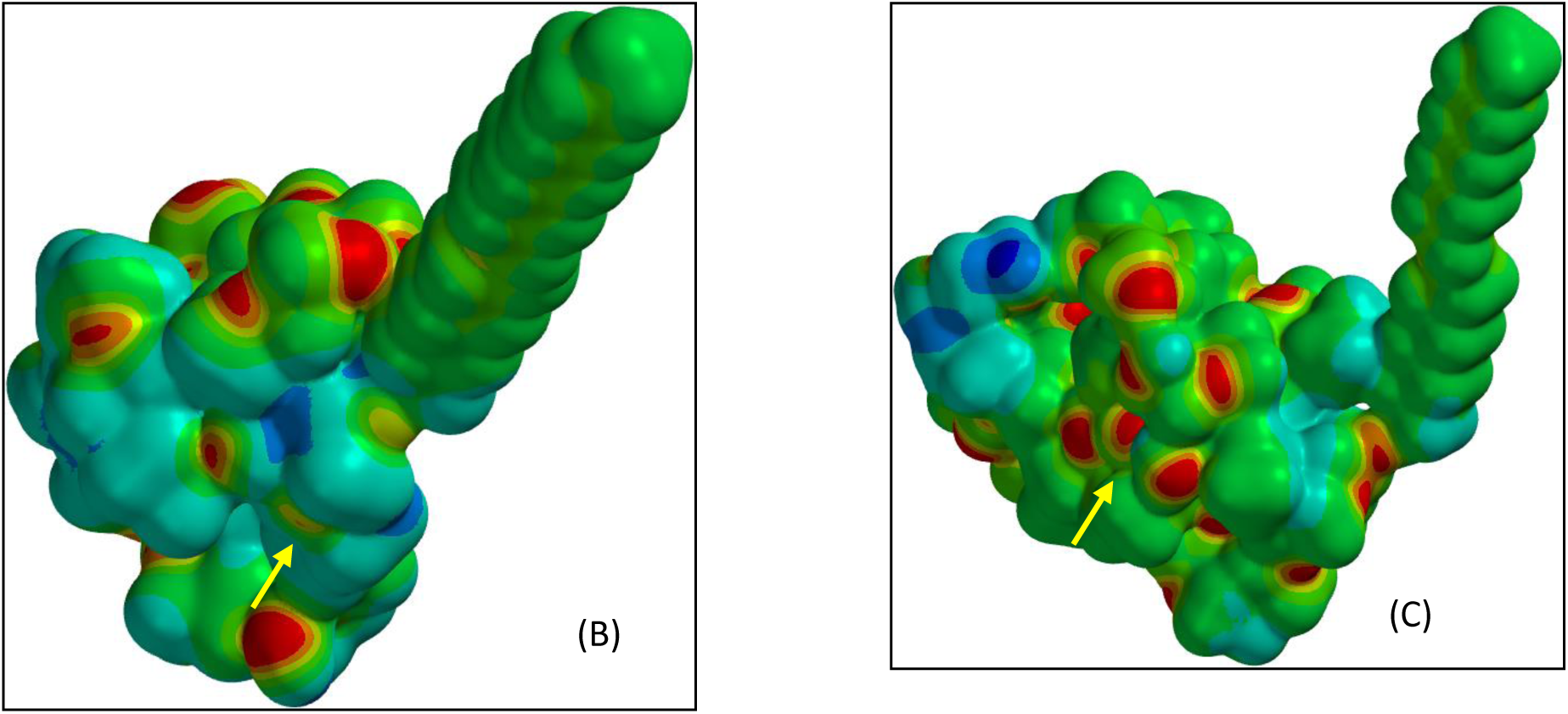
Comparison of Electrostatic Potential Maps (EPM) (A) P-80 molecule without interaction; (B) P-80 molecule interacting with 15 water molecules at positions 1-15; (C) P-80 molecule interacting with 15 water molecules at positions 1-15; and this in turn interacting with 15 ethanol molecules (CH_3_CH_2_OH) at the same sites.

**Figure 31.**
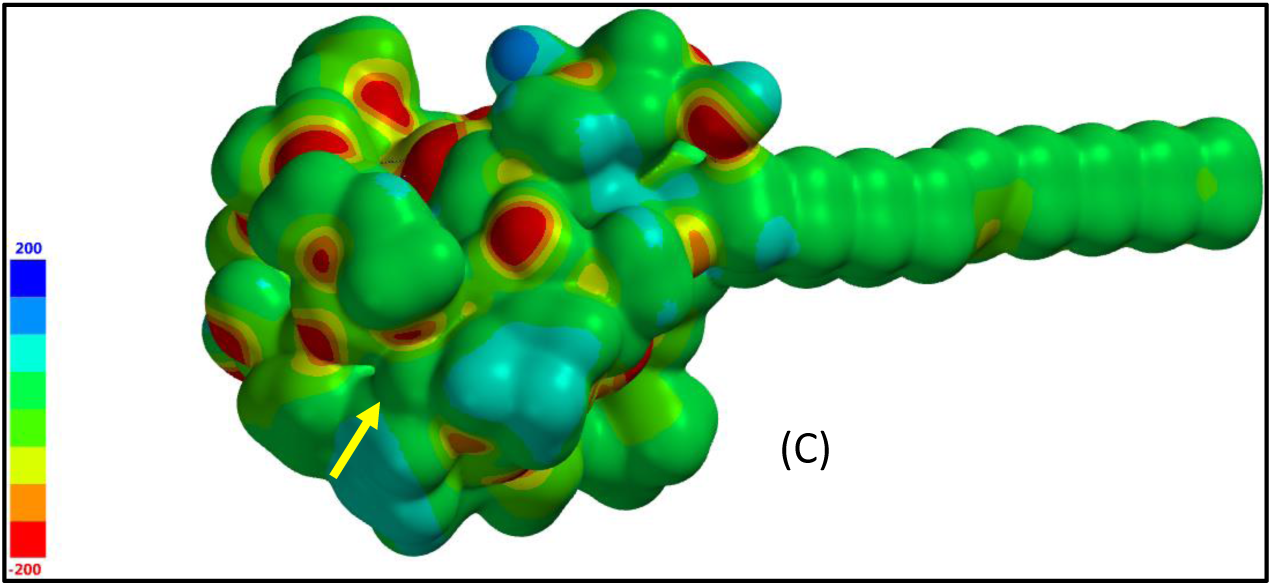
Electrostatic Potential Map (EPM) (C) P-80 molecule interacting with 22 water molecules at positions 1-21; and this in turn interacting with 9 ethanol molecules (CH_3_CH_2_OH) at positions 0-9.

**Figure 32.**
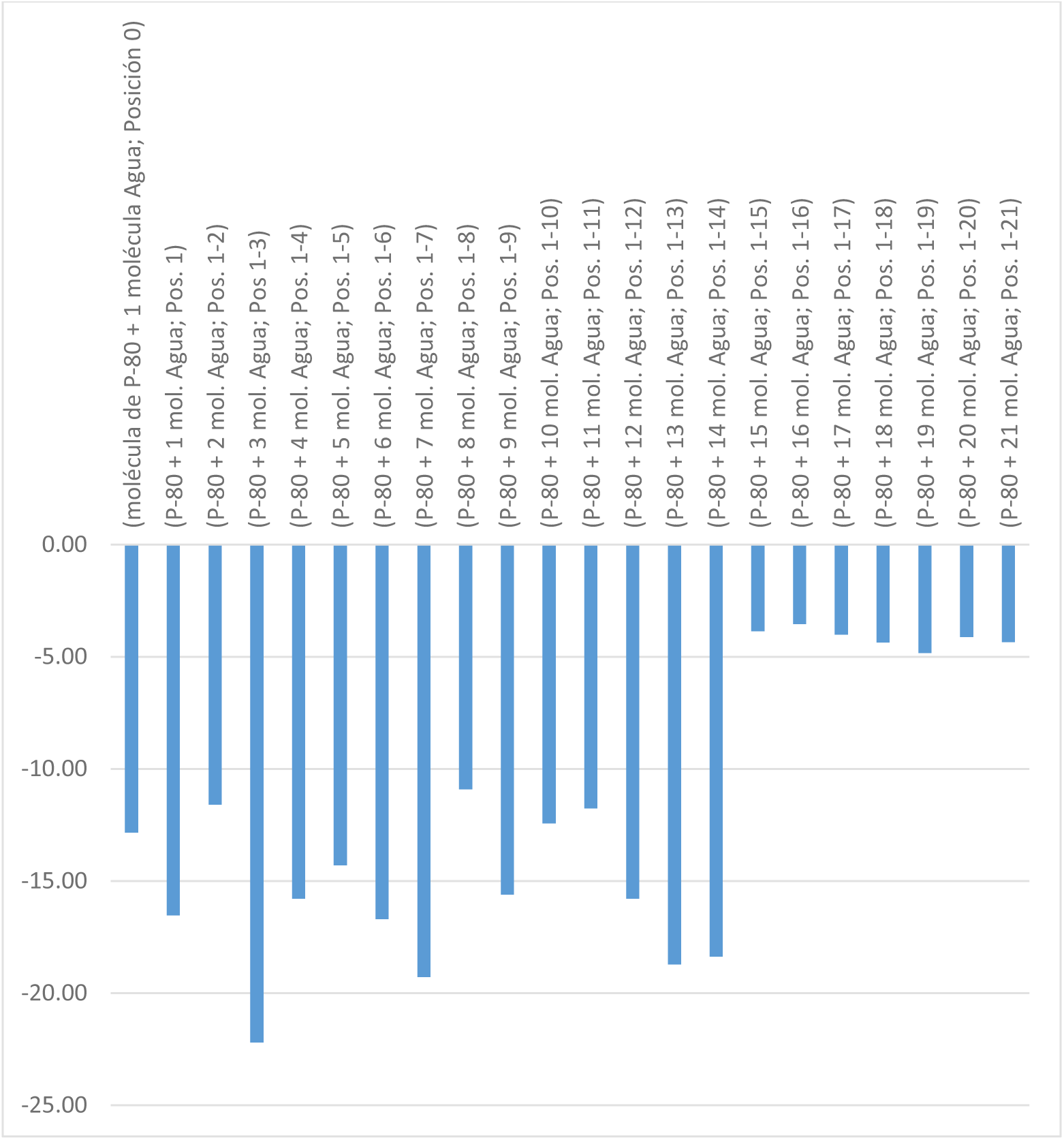
Results of the **sio Index** calculated for each of the molecular forms derived from the interactions of P-80 from 0 to 21 Water molecules, successively (X axis corresponds to the successive substitutions; Y axis refers to the **sio Index** x 10^3^).

**Figure 33.**
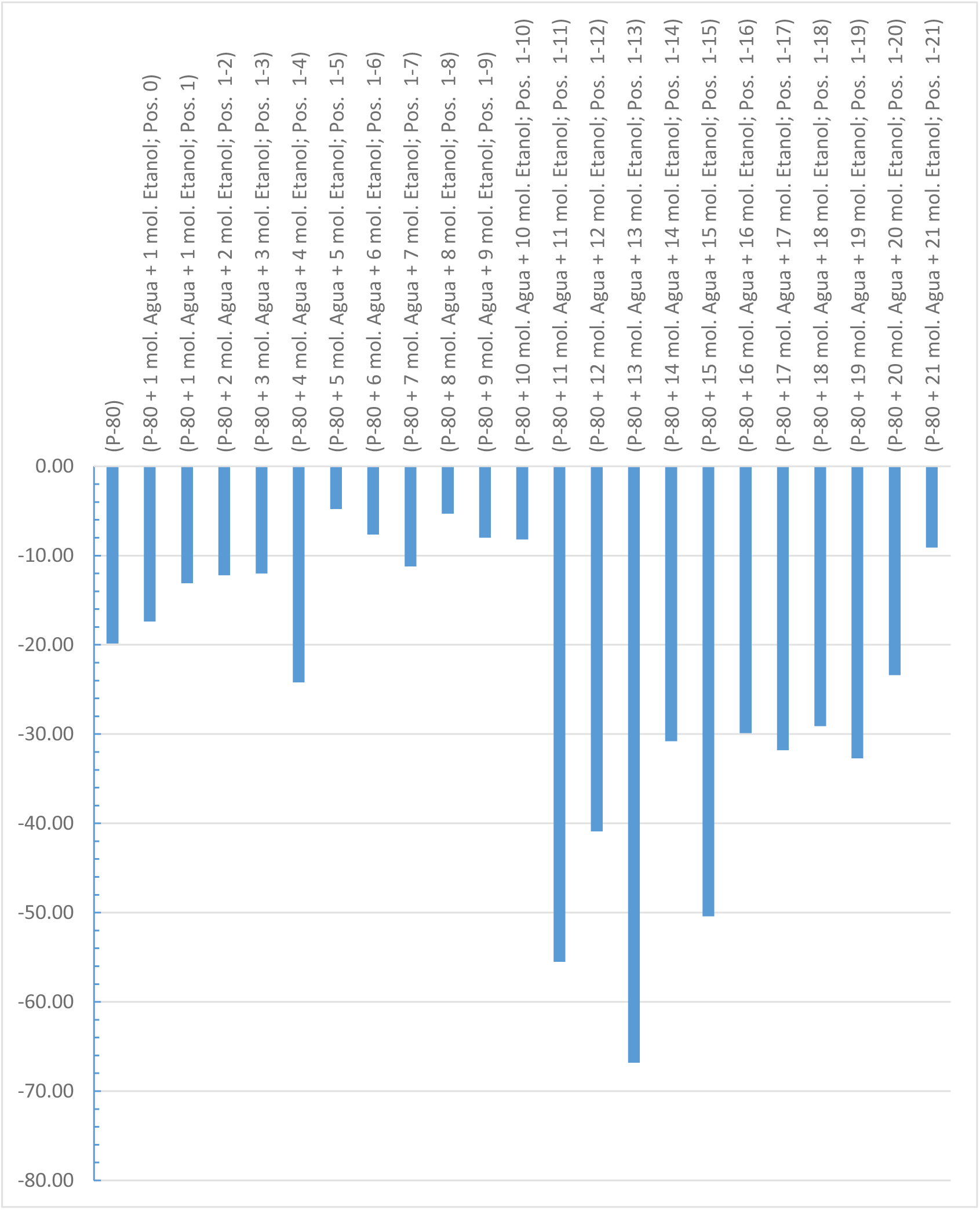
Results of the **sio Index** calculated for each of the molecular forms derived from the interactions with P-80 1 by 1 successively, from positions 0 to 21 with Water and Ethanol molecules, respectively (X axis corresponds to the successive substitutions; Y axis refers to the **sio Index** x 10^3^).

**Figure 34.**
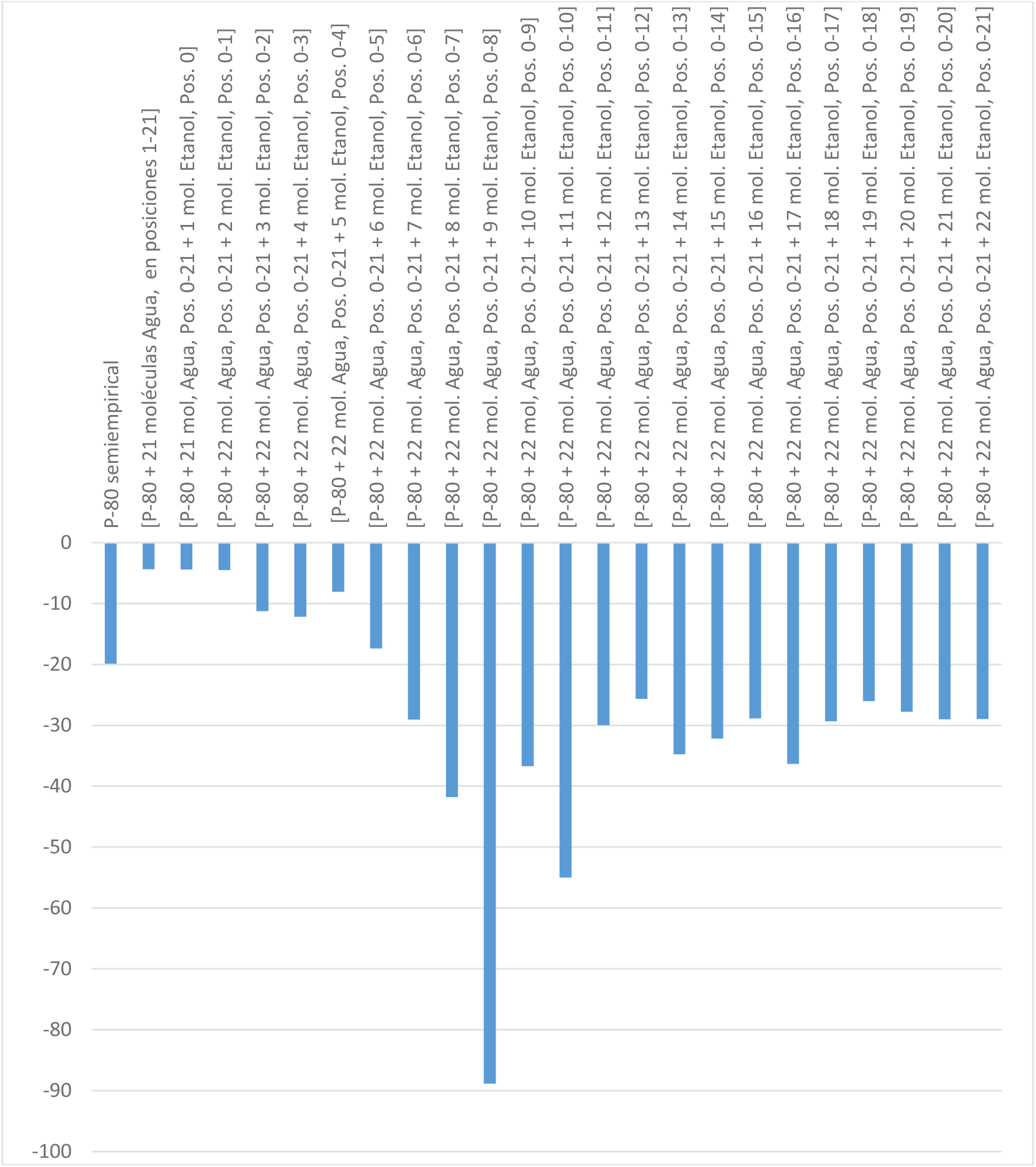
Results of the **sio Index** calculated for each of the molecular forms derived from the interactions of P-80 with positions completely occupied with Water molecules and successively adding from positions 0 to 21 with Ethanol molecules (X axis corresponds to the successive substitutions; Y axis refers to the **sio Index** x 10^3^).

The **sio Index** is relatively low for all molecular forms when simulated with only water (see Table 1, Figure 32). Only the value of 22.20 x 10^-3^ of P-80 substituted with 3 water molecules stands out. The **sio Index** values of the molecular forms with 15 water substitutions and above are lower and similar, compared to the previous molecular forms. In general the values of this series are lower than the values of the series represented in Table 2, Figure 33 and the values of the series represented in Table 3, Figure 34, indicating that the addition of Ethanol to the mixture is important.

The sio Index is relatively low in all molecular forms when simulated by placing 1 water molecule and 1 Ethanol molecule simultaneously, successively up to 10 substitutions (see Table 2, Figure 33). Furthermore, the values in this segment are lower than those in Table 1, Figure 32, however, from 11 substitutions to 21 substitutions the **sio Index** of the derived molecular forms increases significantly. In particular, the molecular forms with 11-15 substitutions have **sio Index** values that are considerably higher than the overall average of the three Tables. This seems to indicate that they are interrelated in some physicochemical way. The role of ethanol in improving and recovering oil is again highlighted in this result. The 13-substitution molecular form is postulated as the optimal form for forming dimers and oligomers.

In Table 3, Figure 34, it can be observed that when the P-80 molecule is saturated with water and Ethanol is added 1 by 1, the sio Index values are low from position 0-4. Again, as in the previous case, From position 5 onwards to the end, the values are relatively high, with an absolute maximum for the molecular form substituted with Ethanol at position 0-8 (**sio Index**=88.85 x 10^-3^). As in the previous case, there seems to be an interrelationship between the molecular forms neighbouring it.

According to these results, the presence of water facilitates to a certain extent the interaction of P-80 with Ethanol and Petroleum, producing its improvement and recovery.

There appears to be a relationship of multiples of 3 and 4 between the maximum values of the “**sio Index**” of Series 2 and Series 3, respectively, in relation to the maximum value of Series 1. The exact values of this relationship are: S2max/S1max=3.009***; S3max/S1max=4.00225***; The *** symbols indicate that there is a periodic repetition of the last three decimal digits. Furthermore, the results as a whole seem to have a tendency to respond to a Fibonacci Series. In previous real experimental results [14] the Oil treated with these Mixtures responded with a greater or lesser improvement of its Density and its °API according to the use of even or odd n-alcohols, respectively, in said Mixture. This result is consistent with the structural periodicity of the Ethoxides, which are repeated even residues (–CH2-CH2-O) – in the polymer chains of P-80. In this case, it is concluded that the Blend would promote the Oil to perform a structural scrutiny. We propose that this structural scrutiny would be done by “Host-Guest Chemistry” in which P-80 is the “Host” and Alcohol is the “Guest”. We propose that the generated nanoparticle [14] is produced by this mechanism.

## Conclusions

In this work it is postulated that there are at least two different extreme and one intermediate forms of interaction between P-80, Ethanol, Water and ions, with Petroleum.The first would be produced at low concentrations of water and low concentrations of ions and the second would be produced at high proportions of water and sodium chloride, both in relation to the other components (Petroleum, P-80 and Ethanol).The intermediate molecular form would be produced between both conditions. The first structure generated would be flatter and with a significant number of P-80 molecules interacting with few water and ethanol molecules. The second structure generated would be more spherical. The intermediate structure would have properties with more evident characteristics of oligomeric association. According to the results of this work, the two forms of the structures generated in the first case [Figure 5 (B), DPM=3.09 and P=154.71; Figure 6 (C), DPM=2.52 and P=176.85] are produced by the interaction of hydrogen bonds of Water and Ethanol with P-80 that resulted in an improvement of the Viscosity, At low proportions of water and ions they should be stabilized as oligomers with a relatively tubular or prolate structure (“Prolate ellipsoid of revolution”) [6]. The second would be more spherical (“Oblate ellipsoid of revolution”)and with a different proportion of the three types of molecules: P-80, Ethanol and Water. Both structures, the prolate and the oblate, would form Oligomers, the first one flatter, with 4 Water molecules and 4 Ethanol molecules associated with each P-80 participating in the Oligomer and the second structure would be more spherical, with 21 Water molecules and 4 to 9 Ethanol molecules associated with each P-80 molecule within this second type of Oligomer [Figure 10 (D), DPM=3.09 and P=38.86]. This second one would be the one that improves the density of the crude oil. An intermediate molecular form between these two and that would improve both the viscosity and the density, is produced in the second series, in which 1 to 1 water and ethanol are added: the ideal molecular form for this would be the one that contains P-80: 13 water molecules: 13 ethanol molecules [see Figure 15 (C) DPM=4.15 and P=38.48].

One question that remains to be resolved is which physicochemical parameters are most relevant to decide which molecular forms are most appropriate to improve and recover oil, for example: if we compare some values of the physicochemical parameters between the P-80 molecule + 3 water molecules (Figure 5(B) and in Table 1, (P-80 semiempirical-3 molH2O (P-80 + 3 mol. Water; Pos. 1-3)) with the P-80 molecule + 22 water molecules + 9 ethanol molecules (Figure 31(C) and Table 3, (0-8) + P-80 semiempirical-21 molH2O+ (0-8) mol ethanol [P-80 + 22 mol. Water; Pos. 0-21 + 9 mol. Ethanol; Pos. 0-8), both with DPM 3.09 Debye, the latter appears to be more stable than the former due to the low energy levels it presents (Minimum Conformational Energy: EMC, Energy:E and sio Index), compared to the former.

It is concluded that the possible impediments to achieving the total improvement of Petroleum as a “Selector” could be explained by the following hypotheses:

1. The molecule that contributes to the desired solution cannot enter the petroleum and that is why the total improvement in Viscosity and Density does not occur.
2. The molecule that contributes to the desired solution does manage to enter the oil, but is expelled due to its stability conditions within it.
3. It may be a problem of oligomeric, flat or spherical conformational structure.

## Acknowledgements

Research Project: ULA Biotechnology INTEVET PDVSA (UBIT, 2006). Project Manager, EDJA.

This work was financed by the Authors. The Authors declares no Competing or Conflict of interests.

## Author Contribution

EDJA and HCG, conceived and designed the UBIP Project.

JGC-M, LEO-G, MAF-R and AYL-R performed simulations.

JGC-M, EDJA, LEO-G and AYL-R performed calculations including the new parameter “sio Index” developed by EDJA.

EDJA, HCG, EAP-C and VJA-G, contribute Real Experiments. JGC-M, EDJA, LEO-G, MAF-R and AYL-R wrote the paper.

